# Combined inactivation of RB and Hippo pathways converts differentiating photoreceptors into eye progenitor cells through derepression of *homothorax*

**DOI:** 10.1101/2023.04.23.537991

**Authors:** Alexandra E. Rader, Battuya Bayarmagnai, Maxim V. Frolov

## Abstract

The RB and Hippo pathways interact to regulate cell proliferation and differentiation. However, their mechanism of interaction is not fully understood. *Drosophila* photoreceptors with inactivated RB and Hippo pathways specify normally but fail to maintain neuronal identity and dedifferentiate. We performed single-cell RNA-sequencing to elucidate the cause of dedifferentiation and the fate of these cells. We find that dedifferentiated cells adopt a progenitor-like fate due to inappropriate activation of the retinal differentiation suppressor *homothorax* (*hth*) by Yki/Sd. This results in activation of the Yki/Hth transcriptional program, driving photoreceptor dedifferentiation. We show that Rbf physically interacts with Yki which, together with the GAGA factor, inhibits *hth* expression. Thus, RB and Hippo pathways cooperate to maintain photoreceptor differentiation by preventing inappropriate expression of *hth* in differentiating photoreceptors. Our work accentuates the importance of both RB and Hippo pathway activity for maintaining the state of terminal differentiation.

## INTRODUCTION

It is well appreciated that the extent of differentiation within a tumor profoundly determines its aggressiveness, dissemination, regenerative potential, and ultimately, its vulnerability to therapy. Tumor cells which adopt stemlike phenotypes possess significant advantage as they proliferate without restriction, evade immune surveillance, and adapt rapidly to therapeutic interventions, ultimately resulting in more aggressive cancer^1,2^. In many cases, transition to a progenitor-like state is coupled with loss of Retinoblastoma (RB) tumor suppressor. RB is necessary for the appropriate specification of many tissues^3–5^ and its loss sensitizes cells to dedifferentiation^6–8^. However, lack of functional RB is not always sufficient for cell fate reversal. Rather, it is hypothesized that dedifferentiation is achieved through aberrant activation of developmental pathways which regulate stemness and renewal of developing tissues^9,10^. In this vein, loss of Hippo pathway inhibition of its effector, YAP, is associated with dedifferentiation in liver^11,12^, pancreatic^13^, kidney^14^, and colorectal^15^ carcinomas, resulting in highly proliferative progenitor-like cells. Despite these observations, the fundamental role of the Hippo pathway in maintaining differentiation of terminally specified cells has not been fully explored. This conspicuous gap in knowledge exists in-part due to limitations in modeling and characterizing YAP-driven dedifferentiation at single-cell resolution. Such experiments require a model for both cell development and subsequent dedifferentiation with minimal genetic heterogeneity and compensation.

The *Drosophila* third instar larval eye imaginal disc is an ideal model for studying cell differentiation *in vivo*. During this developmental stage, the eye tissue is halfway complete with its transition from proliferating eye progenitor cells into mature photoreceptor neurons (R cells). This transition occurs in the morphogenetic furrow (MF), a physical indentation that initiates in the posterior and advances towards the anterior throughout the third instar. This progressive development provides a snapshot of neuronal differentiation over time, where cells that are further posterior from the MF are further differentiated.

The order in which photoreceptors are recruited to form ommatidia is another important aspect of *Drosophila* eye development. Photoreceptor specification begins with the establishment of the R8 photoreceptor which uniquely expresses Senseless (Sens) transcription factor^16^. This founder R8 cell sequentially recruits the remaining R1-7 cells to generate a full ommatidial cluster. As the cluster forms, every R cell eventually expresses the late neuronal marker Elav. Since the R8 cell is absolutely required for recruitment of other R cells, each Elav-positive ommatidium always includes a Sens-positive R8 cell. Surprisingly, combined inactivation of RB and Hippo pathways results in photoreceptor dedifferentiation. This phenotype is manifested through the progressive loss of photoreceptor markers in the posterior and the presence of Elav-positive ommatidia which lack a Sens-positive cell^17^. One of the challenges in this model of dedifferentiation is identifying and characterizing dedifferentiated cells. Thus, the fate of these cells remains unknown.

Another critical gap in knowledge is the understanding of how the RB and Hippo pathways interact to prevent dedifferentiation. E2f1 transcription factor, a downstream effector of the RB pathway, has been shown to cooperate with the *Drosophila* YAP ortholog Yki to promote inappropriate proliferation in the eye posterior^18^. Further, E2f1 was shown to prevent activation of key Yki target genes by disrupting Yki/Sd complexes in Rbf deficient wing discs^19^. However, E2f1 is not required for dedifferentiation in RB and Hippo pathway double mutant photoreceptors^17^ and the role of Yki in this process has not been fully explored. These observations highlight the importance of identifying the dedifferentiation transcription program. However, previous genome wide profiling using total RNA from whole double RB and Hippo mutant eye discs was uninformative in this respect^18^. This is likely due to the extensive cellular heterogeneity of the eye, which prevents extracting the transcriptional signature of dedifferentiating photoreceptors from bulk RNA samples.

To circumvent these challenges, we employed Single-Cell RNA-Sequencing (scRNA-seq) to identify the transcriptional program of dedifferentiating photoreceptors in an unbiased manner. Genetic and scRNA-seq experiments revealed that ectopic activation of Yki in *Rbf* mutant photoreceptors results in aberrant expression of the anterior progenitor gene *homothorax* (*hth*) in the posterior compartment. *hth* upregulation in *yki* expressing tissues is both necessary and sufficient to induce dedifferentiation. Unlike the Yki/Sd pro-proliferative program, Yki/Hth reactivates a particular transcriptional program which imparts photoreceptors with an undifferentiated, progenitor-like phenotype. Interestingly, we find that Yki and Rbf physically interact and that Rbf limits Yki/Sd activation of the *hth* reporter in transcriptional assays. We further show that GAGA factor (GAF), which is known to interact with both Rbf and Yki, limits Yki/Sd activation of *hth* and, like *Rbf*, is required to maintain photoreceptor identity. This suppressive activity by GAF is distinct from its canonical role to enhance Yki dependent transcription of proliferation genes ^20,21^. Thus, our results provide a molecular explanation for how RB and Hippo pathways cooperate to maintain photoreceptor differentiation and identify *hth* as a key target upon which the two pathways converge.

## RESULTS

### Genetic labeling of dedifferentiating *Rbf^120a^wts^x1^* double mutant photoreceptors using Flybow

The conclusion that combined inactivation of RB and Hippo pathways leads to dedifferentiation is based on the progressive loss of photoreceptor markers in clones of *Rbf wts* double mutant cells in the eye disc posterior^22^. One limitation of this analysis is that dedifferentiating cells are identified by the loss of marker and, therefore, the fate of these cells remains unknown. To overcome this caveat, we used elements of the Flybow system^23^ to label photoreceptors by the expression of a *lacZ* reporter. Once activated, this reporter is continually expressed regardless of photoreceptor differentiation status (Figure 1A). Importantly, Flybow utilizes a modified inducible mFlp5/mFRT71 to turn on *lacZ* gene expression and is compatible with the conventional ey-Flp/FRT system, thus, allowing clones of *Rbf wts* double mutant cells to be simultaneously generated.

**Figure 1.**
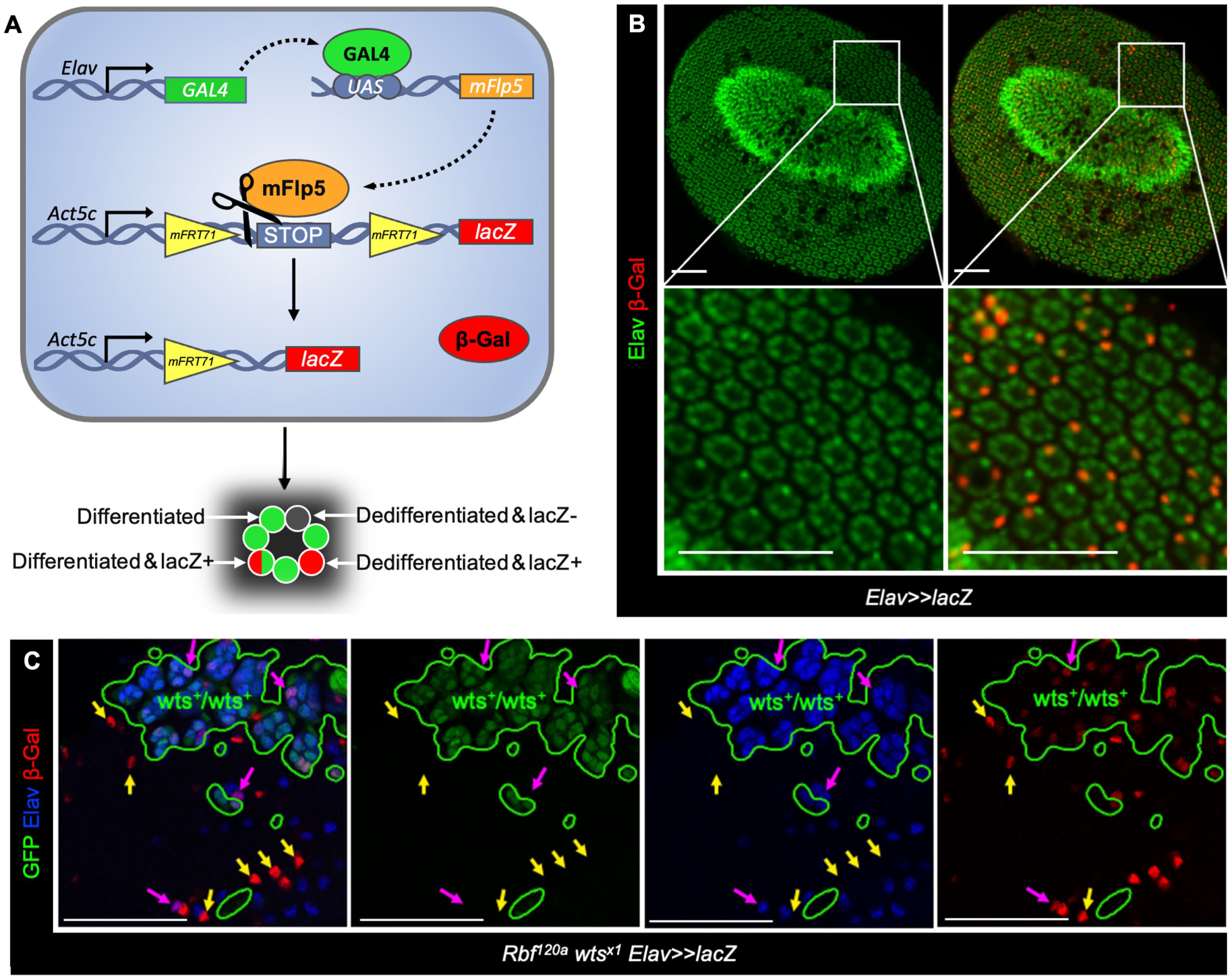
Genetic labeling to visualize dedifferentiating *Rbf wts* double mutant photoreceptors. (A) A schematic of the Flybow based system to label photoreceptors. Elav-Gal4 drives expression of *uas-mFlp5* to allow the persistent *lacZ* expression regardless of the activity of the Elav promoter. (B) 48 hr APF pupal eye from a *+/+;uas-mFlp5, Act5c<stop<lacZ/+;Elav-Gal4/+* fly stained for Elav (green) and β-Gal (red). (C) 48 hr APF pupal eye of *Rbf^120a^, ey-Flp/Y;uas-mFlp5, Act5c<stop<lacZ/+;Elav-Gal4, FRT82B, GFP /FRT82B, wts^x1^* fly stained for Elav (blue) and β-Gal (red). Wild type cells are labeled by GFP, homozygous *wts^x1^* mutant cells are GFP-negative. Elav-negative, β-Gal-positive cells visualize *Rbf wts* double mutant photoreceptors that fail to maintain their neuronal identity. Scale bars are 50 μm.

We began by determining the efficiency of the photoreceptor labeling system in wildtype eyes. We combined the *act≥stop≥lacZ* (≥ indicates mFRT71) construct with *uas-mFLP71* and used *Elav-GAL4* to express *mFLP71* in differentiating photoreceptors. Pupal eyes were dissected at 48 hrs APF, stained with Elav and β-Gal and the number of β-Gal positive cells per ommatidium was determined (Figure 1B). As expected, every β-Gal-positive cell was also positive for Elav, indicating that photoreceptors are indeed specifically labeled. Conversely, not every photoreceptor expressed β-Gal. On average, there were two β-Gal-positive cells per ommatidial cluster, with ommatidia containing between zero and three β-Gal-expressing cells, indicating that less than half of cells in the cluster were labeled by 48 hrs APF.

Next, we induced clones of *Rbf wts* double mutant cells with *ey-FLP* and used the aforementioned transgenes to assess β-Gal positive photoreceptors in pupal eyes at 48 hrs APF. In the wildtype tissue, marked by the presence of GFP, every β-Gal positive cell maintained expression of Elav (Figure 1C). In contrast, there were multiple examples of β-Gal positive, Elav negative cells within *Rbf wts* double mutant clones (marked by yellow arrows in Figure 1C). Since the presence of β-Gal indicates past expression of Elav, this observation suggests that *Rbf wts* double mutant cells initially express Elav but lose expression over time, a hallmark of dedifferentiation. We note, however, that the number of dedifferentiated photoreceptors is likely severely underestimated due to the relative inefficiency of the labeling system (Figure 1B). In addition, cells that do not activate *lacZ* prior to losing Elav expression and dedifferentiating remain unlabeled. Given that the onset of dedifferentiation appears a few columns posterior to the MF^22^, this will significantly impact the number of β-Gal positive dedifferentiated cells. Nevertheless, the appearance of β-Gal positive, Elav negative cells unambiguously demonstrates that *Rbf wts* double mutant photoreceptors are not eliminated and indeed dedifferentiate into an unknown cell type.

### Single-cell RNA-Sequencing reveals upregulation of *hth* in dedifferentiating *Rbf wts* double mutant cells

The relative inefficiency of Flybow to label dedifferentiating photoreceptors essentially precluded a conventional marker analysis to characterize these cells and prompted us to seek alternative approaches. Single-cell RNA sequencing (scRNA-Seq) technology has proven to be highly informative for dissecting cell heterogeneity and identifying unique cell populations during development and in disease^24–28^. The *Drosophila* larval eye imaginal disc was shown to be highly amenable to scRNA-seq profiling, allowing for the identification of all known cell types, as well as mutant specific cell populations^29,30^. Therefore, we employed scRNA-seq to identify and characterize *Rbf wts* double mutant dedifferentiating photoreceptors in an unbiased manner.

Third instar larval eye discs from wildtype and *Rbf* mutant animals and eye discs carrying clones of *wts* single and *Rbf wts* double mutant cells were dissected at the optic stalk and antennal border, dissociated into single cell suspension, and processed using 10X Chromium. The sequencing data from two replicates of wildtype and *Rbf* mutant eye discs and single replicates of *wts* single and *Rbf wts* mutant mosaic eye discs were processed in Seurat 3. After filtering poor quality cells, a total of 4,232 wildtype, 2,643 *Rbf* mutant, 2,158 *wts* mutant, and 2,643 *Rbf wts* double mutant cells, with an average of 2,104 genes per cell were recovered. The Uniform Manifold Approximation and Projection (UMAP) dimensionality reduction algorithm was used to visualize cell populations across all four genotypes. Cell populations were assigned using markers that are specific for each cell population (Table S1). Comparison with our published eye disc cell atlas generated by Drop-seq^29^ revealed that all previously identified cell populations were represented in the new dataset (Figure 2A). We followed the nomenclature of the published cell atlas of the larval eye disc^29^ and applied the same names to cell populations identified here. Using population specific markers we identified two clusters of asynchronously proliferating cells anterior to the morphogenetic furrow, UND and PPN, early and late photoreceptors (EPR and LPR), interommatidial cells (INT), cells from the morphogenetic furrow (MF), and others that were named as in Ariss et al. 2018^29^ (Figure 2A). Importantly, an *Rbf* mutant specific cell population (APOP) characterized by expression of apoptotic and glycolytic genes was readily detected in the *Rbf* mutant sample. Overall, cells from each genotype contributed to every cell population and there were no significant changes in gene expression within clusters between the genotypes. We noticed however, that *Rbf* and *wts* single mutant cells of the pre-proneural (PPN) population clustered discretely from the wildtype (Figure S1A). Analysis of differentially expressed genes in the *Rbf* sample revealed that this clustering was driven by upregulation in cell cycle genes while the PPN *wts* mutant cells upregulated expression of the known Yki target gene *ex* and genes associated with cuticle development. However, both *Rbf* and *wts* PPN cells still expressed the classical PPN markers, including *h* and *dac*^29^ and were therefore merged with their wildtype counterparts in downstream analyses (Figure S1A). We concluded that the newly generated scRNA-seq datasets match well with the previously published cell atlas and are highly reproducible across different technical platforms.

**Figure 2.**
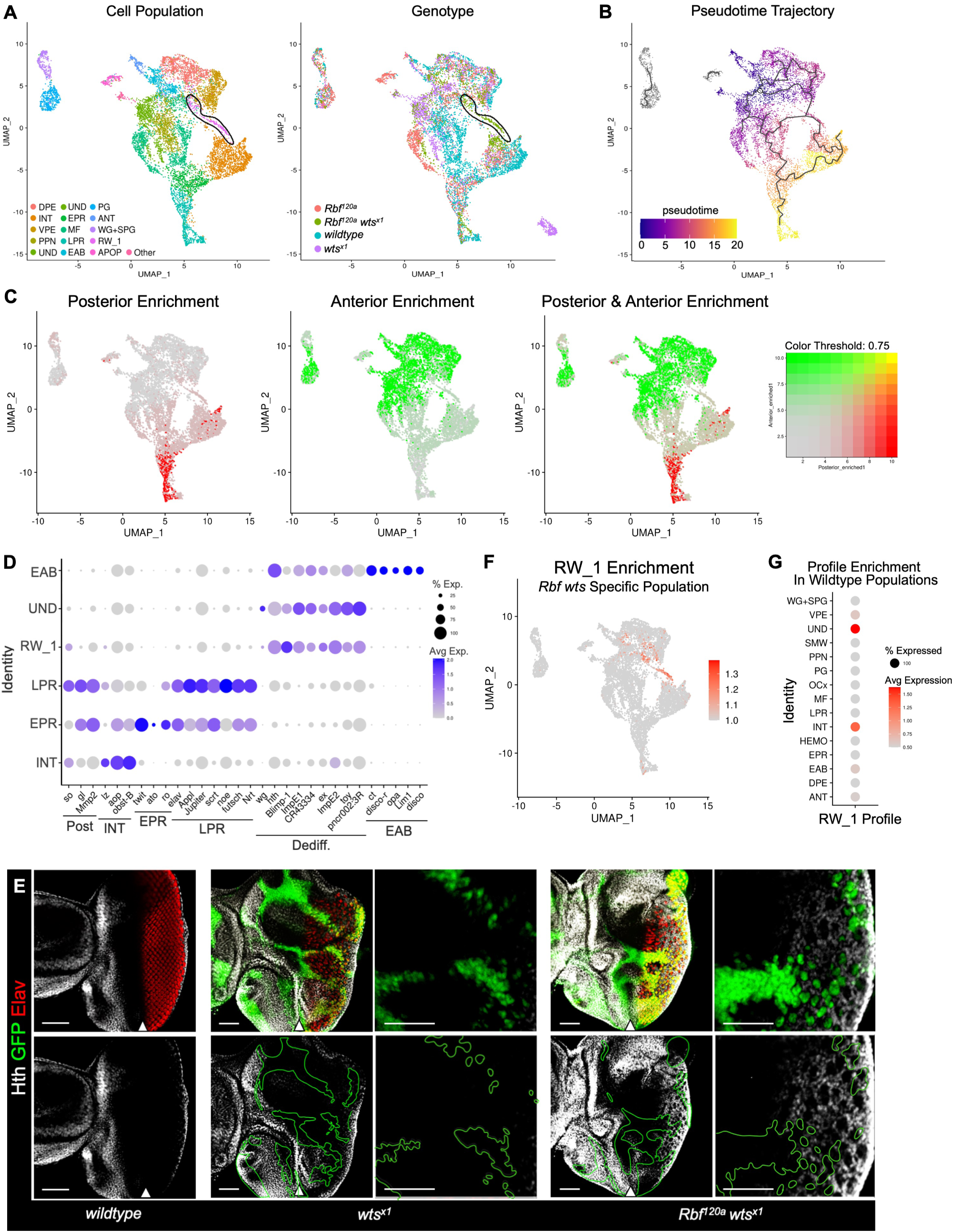
Mosaic *Rbf wts* double mutant eye discs inappropriately expresses *hth* in the posterior. (A) An annotated UMAP of cells by population (left) and sample genotype (right) in *w^1118^*, *Rbf^120a^* mutant, clonal *wts^x1^* mutant, and *Rbf^120a^ wts^x1^* mosaic double mutant tissue. Previously described cell types were identified with the addition of a novel double mutant specific population labeled RW_1 and circled in black. (B) Trajectory analysis performed using Monocle3 depicting cell differentiation states in pseudotime. RW_1 cells arise from the posterior compartment but reverse in pseudotime. Predicted specification from the root node (white circle) is shown with gray lines. (C) Feature plots visualizing enrichment for transcriptional profiles characteristic for anterior and posterior cells across cell populations. (D) A Dot Plot of markers for posterior cells (Post), interommatidial cells (INT), early photoreceptor neurons (EPR), late photoreceptor neurons (LPR), RW_1 (Dediff), undifferentiated cells (UND), and the eye-antennal border (EAB). Expression is shown in corresponding cell populations. The percentage of cells within a population expressing a given gene is shown through dot scale and level of expression is reflected by color intensity. (E) Confocal images of third instar larval eyes of *ey-Flp/Y; FRT82B GFP /FRT82B wts^x1^* and *Rbf^120a^ ey-Flp/Y; FRT82B GFP /FRT82B wts^x1^* animals stained for Elav (red) and Hth (white). GFP-positive tissue is homozygous *wts* wildtype, GFP-negative is homozygous *wts^x1^*mutant. Hth is restricted to the anterior compartment and posterior margin in *wts^x1^* mutant tissue while *Rbf^120a^wts^x1^* double mutant tissue ectopically express *Hth* in the posterior. Scale bars are 50 μm. (F)A feature plot displaying expression of the RW_1 transcriptional profile. The RW_1 Enrichment profile accurately and specifically captures RW_1 cells. (G) Dot Plot of the RW_1 transcription profile on wildtype cell populations. RW_1 most closely resembles UND and INT populations.

Since dedifferentiation only occurs in *Rbf wts* double mutant cells, we hypothesized that these dedifferentiating cells may comprise a distinct cell population. Indeed, UMAP plot showed that the RW_1 cluster consisted entirely of *Rbf wts* double mutant cells, while other populations contained cells of all genotypes (Figure 2A). The name RW_1 was used to reflect that this cell population is unique for *<u>R</u>bf <u>w</u>ts* (RW) double mutant cells. To characterize temporal order of the RW_1 cluster, we performed trajectory analysis using the Monocle 3 package which visualizes the position of each cell along the pseudotime differentiation axis^31^. Uncommitted cells from the furthest anterior region, the EAB, were used as the root node and an indicator of the earliest cell population in pseudotime (Figure 2B). As expected, this analysis revealed the normal progression of photoreceptor differentiation in the eye from EAB, UND, and PPN progenitor cells in the anterior compartment to INT, EPR and LPR in the posterior compartment, as well as a bifurcation between INT and EPR as cells exit the MF (Figure 2B). Intriguingly, the EPR connects in pseudotime with RW_1 through INT, while RW_1 is also connected with the anterior populations PPN, UND, and EAB. Furthermore, RW_1 cells are positioned relatively early along the pseudotime neuronal differentiation axis.

As RW_1 branches between both anterior and posterior populations, we next assessed from which compartment RW_1 originates. To do this, we performed differential expression analysis on anterior and posterior compartment populations to generate a transcriptional profile characterizing either compartment (Table S2). Feature plots for enrichment in each transcriptional profile reveal that these gene modules accurately capture their respective cell populations; the MF, EPR, LPR, and INT are enriched for the posterior transcriptional profile, while the EAB, PPN, and UND upregulate genes associated with the anterior compartment (Figure 2C). But most interestingly, RW_1 was solely enriched for the posterior transcriptional program (Figure 2C), indicating that RW_1 is in fact a posterior cell population. Taken together, this suggests that RW_1 is comprised of posterior *Rbf^120a^ wts^x1^* double mutant cells which experience a reduction in pseudotime.

In order to identify possible markers for RW_1 cells, we generated a Dot Plot to compare the expression of top markers between RW_1 and clusters which were linked in trajectory analysis, including: the EAB, UND, INT, EPR and LPR. One of the top markers for RW_1 is the homeodomain transcription factor and Yki binding partner *homothorax* (*hth*), its target gene *bantam*^32^, as well as several nuclear hormone receptors (NHRs) which are known to respond to *hth* overexpression including *Blimp-1* and *ImpE1*^33^ (Figure 2D). This is significant since Hth is a known suppressor of photoreceptor differentiation and is expressed in eye progenitor cells of the anterior compartment^34,35^,. Accordingly, *hth*, *bantam*, *Blimp-1* and *ImpE1* are also highly expressed in the anterior UND and EAB clusters. However, in addition to enrichment for posterior genes (Figure 2C), RW_1 does not express other EAB top markers such as *Lim1* or *disco*. This suggests that RW_1 is distinct from the EAB.

Next, we examined the expression of markers for posterior cell populations in the *Rbf wts* specific RW_1 population (Figure 2D). RW_1 expresses low levels of the posterior markers *sin oculus* (*so*) and, to a lesser extent, *Mmp2*. But, it fails to express other posterior markers such as *glass* (*gl*) which is present at high levels in INT, EPR and LPR (Figure 2D). RW_1 was also clearly distinct from posterior interommatidial cells since the RW_1 cells do not express the INT marker *lozenge* (*lz)*. Accordingly, RW_1 does not express any neuronal markers of EPR or LPR such as *twit, Elav,* or *futsch*. These results suggest that RW_1 is a distinct cluster with some features of posterior and anterior cells. Notably, a known inhibitor of differentiation *hth* and its targets are highly expressed in RW_1.

Given that *hth* is highly expressed in RW_1 cells, we examined the pattern of Hth expression in *Rbf wts* double mutant mosaic eye discs to spatially localize the RW_1 cluster. The eye discs containing clones of *wts* single mutant and *Rbf wts* double mutant cells were stained with Hth antibodies and visualized by immunofluorescence. In the wildtype eye disc, Hth expression is restricted to anterior undifferentiated cells and the posterior margin (Figure 2E). In contrast, Hth was inappropriately expressed in *Rbf wts* double mutant cells of the posterior compartment where photoreceptors undergo dedifferentiation (Figure 2E). Importantly, ectopic *hth* expression in the eye posterior was specific to *Rbf wts* double mutant cells, as it was absent in posterior clones of *wts* mutant cells (Figure 2E). Further, fluorescence quantification confirms that only double mutant tissue significantly upregulates Hth (Figure S1B). These results reveal that *Rbf wts* double mutant cells inappropriately upregulate *hth* in the posterior compartment.

Having confirmed *in vivo* that RW_1 consists of posterior *Rbf wts* double mutant cells which ectopically express the anterior differentiation suppressor *hth*, we next assessed whether RW_1 shared further commonality with undifferentiated cell populations. In other words, do RW_1 cells adopt an anterior-like expression profile? As previously described, we performed differential expression analysis on RW_1 to produce a transcriptional profile which is characteristic of RW_1 cells (Table S3). Feature plot for enrichment of this profile suggests that this signature is unique to and characteristic of RW_1 cells (Figure 2F). We then calculated enrichment of the RW_1 expression profile in wildtype cells alone and generated a Dot Plot to observe which wildtype populations most closely resemble *Rbf wts* double mutant cells (Figure 2G). This Dot Plot showed that RW_1 was transcriptionally similar to both posterior INT cells and anterior UND cells, suggesting that double mutant posterior tissues which ectopically express *hth* adopt an anterior-like features.

### Ectopic expression of constitutively activated Yki in *Rbf* mutant eye discs induces photoreceptor dedifferentiation

The results described above led us to hypothesize that RW_1 consists of dedifferentiating *Rbf wts* double mutant photoreceptors and that *hth* is a one of the top markers of these cells. This is intriguing because *hth* is normally restricted to progenitor cells of the eye disc, anterior to the MF, where its role is to inhibit photoreceptor differentiation^34,35^. Therefore, inappropriate activation of *hth* in the posterior compartment of *Rbf wts* double mutant tissues may explain why these photoreceptors dedifferentiate. To confirm that combined inactivation of Hippo and RB pathways leads to ectopic expression of *hth*, we mimicked Hippo pathway inactivation using a constitutively active Hippo effector, Yki^S111A,S168A,S250A^ (Yki^CA^), that is insensitive to Wts regulation^36^.

*Yki^CA^* was overexpressed in the posterior compartment of wildtype and *Rbf* mutant eye discs using the *GMR-Gal4* driver and expression of neuronal markers Sens and Elav was determined by immunofluorescence. Sens is uniquely expressed in the founder R8 cell while Elav is expressed in all photoreceptors of each ommatidia. Therefore, in the wildtype eye disc, each cluster of Elav positive cells contains a Sens positive cell (Figure 3A). It has been previously demonstrated that dedifferentiation of *Rbf wts* double mutant photoreceptors is visualized by the progressive loss of Elav positive cells and the presence of ommatidia without a Sens positive cell^22^. Overexpression of *yki^CA^* results in extensive expansion of interommatidial cells that disrupts ommatidial organization, however, every ommatidium contains a Sens positive cell (Figure 3A). In contrast, *yki^CA^* overexpression in *Rbf* mutant eye discs severely decreased the number of Elav positive cells toward the posterior of the eye disc and there were numerous examples of Elav positive clusters which lack a Sens positive cell, a hallmark of dedifferentiation (white arrows in Figure 3A). To quantify the extent of dedifferentiation, we counted the number of Elav positive clusters lacking a Sens positive cell using ZEN software (see Materials and Methods). In wildtype retina, 99% of ommatidia contained a Sens expressing cell (Figure 3B). Despite severe disruption in spacing between ommatidia due to proliferation of interommatidial cells in *GMR>yki^CA^*, a Sens positive cell was found in 97% of Elav positive clusters. However, in *Rbf GMR>yki^CA^* eye discs, in addition to severe loss of Elav positive cells, a Sens positive cell was missing in 26% of Elav positive clusters. Additionally, we noticed multiple examples of clusters bearing duplicated R8 cells (circled in Figure 3A), indicating that Yki prevents cell cycle exit of differentiated *Rbf* mutant photoreceptors. This is in agreement with the cell cycle defects previously described in the *Rbf wts* double mutant^17^.

**Figure 3.**
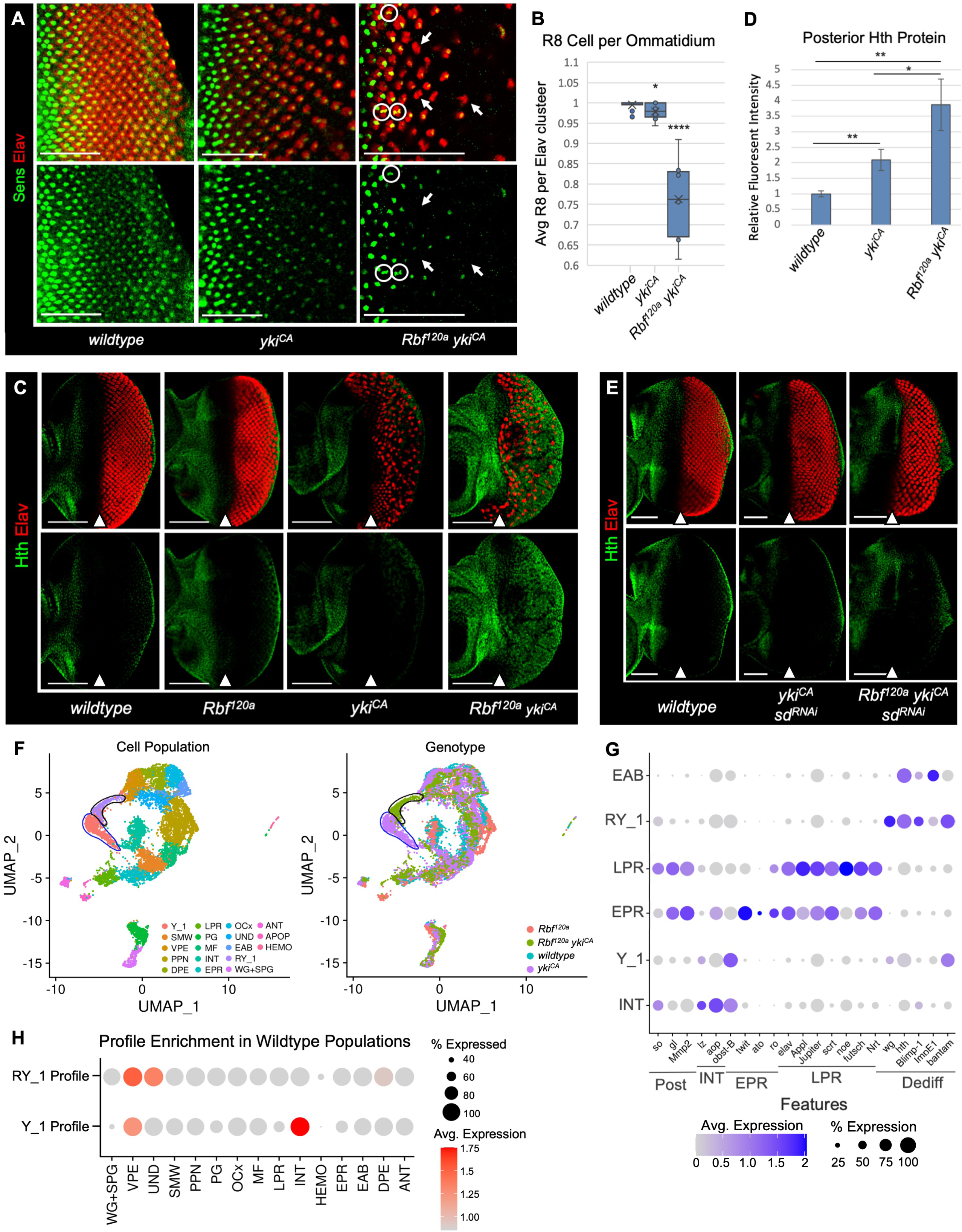
Ectopic *Yki^CA^* induces Hth expression and photoreceptor dedifferentiation in *Rbf* mutant eye discs. (A) Confocal images of eye discs from *w^1118^; GMR-Gal4/+; uas-yki^CA^/+,* and *Rbf^120a^/Y; GMR-Gal4/+; uas-yki^CA^/+* animals stained for Senseless (green) and Elav (red) to visualize R8 cells and all photoreceptors accordingly. Photoreceptor dedifferentiation is visualized by the presence of Sens-negative, Elav-positive ommatidial clusters (white arrows). *Rbf^120a^ GMR>yki^CA^* photoreceptors fail to exit the cell cycle resulting in duplicate R8 cells (white circles). Images are cropped at the morphogenetic furrow. Scale bars are 50 μm. (B) A box-and-whisker plot displaying the ratio of Sens-positive R8 cells per Elav-positive ommatidial cluster. Ommatidia and R8 cell numbers were identified using ZEN software (N=10 eye discs per genotype). The extent of dedifferentiation is scored when there is a significant reduction in the number of ommatidial clusters with an obligatory R8 founder cell. (C) Confocal images of wildtype, *Rbf^120a^, GMR>yki^CA^*, and *Rbf^120a^ GMR>yki^CA^* eye discs stained for Elav (red) and Hth (green). Hth is normally restricted to the anterior compartment and posterior margin in *wildtype*, *Rbf^120a^* mutant, and *GMR>yki^CA^* expressing tissues. Expression of *yki^CA^* in *Rbf^120a^*mutant eye discs results in aberrant Hth protein in the posterior compartment. The MF is labeled with a white arrow. Scale bars are 100 μm. (D) A bar graph displaying Hth protein levels in the posterior compartment normalized to the anterior compartment of each tissue (N=10 eye discs per genotype). There is a significant increase in Hth protein in the posterior compartment of *Rbf^120a^* mutant *yki^CA^* expressing tissue compared to *w^1118^* and *yki^CA^* expressing cells with wildtype *Rbf*. (E) Wildtype, *+/Y;GMR-Gal4/uas-Sd^RNAi^; uas-yki^CA^/+*, and *Rbf^120a^/Y; GMR-Gal4/uas-Sd^RNAi^; uas-yki^CA^/+* eye discs stained for Elav (red) and Hth (green). Knockdown of Sd limits Yki-driven interommatidial expansion and ectopic Hth expression in the posterior compartment. The position of the MF is shown by a white arrow. Scale bars are 50 μm. (F) An annotated UMAP of cells by population (left) and genotype-of-origin (right) in *w^1118^*, *Rbf^120a^*, *GMR>yki^CA^*, and *Rbf^120a^ GMR>yki^CA^*samples. An *Rbf^120a^ GMR>yki^CA^* specific population, designated RY_1, is circled in black and the *GMR>yki^CA^* specific population Y_1 is circled in blue. (G) A Dot Plot of markers for posterior cells (Post), interommatidial cells (INT), early photoreceptor neurons (EPR), late photoreceptor neurons (LPR), and RW_1 (Dediff). Expression is shown in corresponding cell populations, Y_1, and RY_1. Like RW_1, RY_1 upregulates *hth* and its target genes. The percentage of cells within a population expressing a given gene is shown through dot scale and level of expression is reflected by color intensity. (H) Dot Plot of the Y_1 and RY_1 transcription profiles on wildtype cell populations. *GMR>yki^CA^* expressing cells upregulate genes associated with the VPE while *Rbf^120a^ GMR>yki^CA^* cells express an additional UND-like signature.

Having established photoreceptor dedifferentiation in *Rbf GMR>yki^CA^* eye discs, we next asked whether it was accompanied by the presence of ectopic Hth. The expression of Elav and Hth in wildtype, *Rbf^120a^*, *GMR>yki^CA^*, and *Rbf^120a^ GMR>yki^CA^* eye discs was determined by immunofluorescence. As expected, Hth was localized to the anterior compartment and posterior margin in wildtype and *Rbf* mutant eye discs (Figure 3C). In *GMR>yki^CA^,* Hth remained restricted to the anterior compartment and the posterior margin as in wildtype. Strikingly, the *Rbf^120a^ GMR>yki^CA^* eye disc showed a strong *hth* expression immediately posterior to the MF at the onset of *yki^CA^* expression (Figure 3C) which was significantly higher than in other genotypes (Figure 3D).

Yki lacks DNA-binding activity. In the posterior eye, Yki is recruited to DNA by Scalloped (Sd), a TEAD transcription factor, to execute Hippo transcriptional program^37,38^. Therefore, we asked whether inappropriate expression of Hth in the posterior compartment of *Rbf GMR>yki^CA^* eye discs is dependent on Sd. We used RNAi to deplete Sd in *Rbf GMR>yki^CA^*tissues and examined Hth expression by immunofluorescence. As has been previously shown, Sd knockdown by RNAi using *GMR-Gal4* blocks Yki driven inappropriate proliferation as evident by normal spacing between ommatidia (Figure 3E). Notably, depletion of Sd fully prevents inappropriate expression of *hth* in *Rbf GMR>yki^CA^ sd^RNAi^*. This suggests that Yki induces Hth in *Rbf* mutant eye discs in a Sd-dependent manner.

To further characterize dedifferentiating photoreceptors in *Rbf GMR>yki^CA^* and compare with that of mosaic *Rbf wts* double mutant eyes, we performed scRNA-seq using dissected eye discs of *GMR>yki^CA^*and *Rbf GMR>yki^CA^* as described above. After filtering poor quality cells, we retained 4,116 of *GMR>yki^CA^* and 1,798 of *Rbf GMR>yki^CA^* cells (Table S4). We then combined these datasets with that of wildtype and *Rbf* mutant eye discs generated above and performed UMAP analysis to visualize cell populations across all four genotypes (Figure 3F). As expected, *Rbf GMR>yki^CA^* and *GMR>yki^CA^* contributed to every cell population of the wildtype sample. To identify a cluster of dedifferentiating photoreceptors, we used the same approach as for the *Rbf wts* dataset, described above, and isolated a cell population, RY_1, consisting exclusively of *Rbf GMR>yki^CA^* cells (black outline in Figure 3F). The name RY_1 was used to reflect that it is specific for the *<u>R</u>bf <u>y</u>ki* genotype. We then generated a Dot Plot visualizing the expression of EPR, LPR, and RW_1 dedifferentiation markers in the RY_1 cluster. Strikingly, the expression pattern in RY_1 was very similar to that of the *Rbf wts* specific population RW_1; both clusters highly expressed *hth* and nuclear hormone receptors at levels akin to that of in EAB (Figure 3G). We also note that cluster Y_1 contained only *GMR>yki^CA^* cells (blue outline Figure 3F), though Dot Plot shows that Y_1 maintains expression of interommatidial markers, indicating that Y_1 represents inappropriately proliferating interommatidial cells. This aligns with the canonical function of Yki: preventing cell cycle exit in quiescent interommatidial cells.

Finally, we applied a similar approach as with RW_1 to assess the identities of the *Rbf GMR>yki^CA^*specific cell population RY_1, which dedifferentiate, and the *GMR>yki^CA^*specific cell population Y_1, which maintain specification. We performed differential expression analyses on RY_1 and Y_1 to generate profiles which were specifically characteristic of each population (Figure S2A-B, Table S5-6). We then projected these profiles onto wildtype cell populations to identify which normal cell types are most similar to RY_1 and Y_1 cell populations. As shown in Figure 3H, Y_1 cells shared an expression profile similar to that of the VPE and INT. This similarity to peripodial membrane cells of the VPE is in line with previous work which suggests that Yki is primarily active in cells of the peripodial membrane^39^. Importantly, the Y_1 transcriptional profile most closely resembled wild type interommatidial cells INT, but showed no similarity to anterior cell populations. However, the dedifferentiating RY_1 transcriptional profile was strikingly distinct from Y_1. In addition to a peripodial-like signature, the RY_1 profile was highly similar to that of the anterior UND population (Figure 3H). Thus, only in an *Rbf* mutant background, *yki^CA^* expression shifts the transcriptional signature of posterior cells towards anterior undifferentiating progenitor cells. This is in concordance that photoreceptor dedifferentiation occurs only in *Rbf GMR>yki^CA^* but not in *GMR>yki^CA^*eyes. Thus, as with RW_1, posterior RY_1 cells adopt an undifferentiated progenitor-like phenotype.

Therefore, we concluded that combined inactivation of RB and Hippo pathways in mosaic *Rbf wts* double mutant eye discs or in *Rbf GMR>yki^CA^* mutant eye discs results in inappropriate expression of *hth* posterior to the MF and photoreceptor dedifferentiation. scRNA-seq profiling confirmed that these dedifferentiating photoreceptors share similar transcriptional programs which feature *hth* as a top marker.

### *hth* is necessary and sufficient to trigger photoreceptor dedifferentiation

Single cell profiling suggests that a major shared feature of dedifferentiating photoreceptors in either mosaic *Rbf wts* double mutant or in *Rbf GMR> yki^CA^*eye discs is the expression of *hth*. Immunofluorescence experiments confirm that Hth is ectopically expressed in the eye posterior at the point where mutant photoreceptors undergo dedifferentiation. Given the known role of *hth* in inhibiting differentiation of eye progenitor cells, we asked whether inappropriate expression of *hth* in photoreceptors with inactivated RB and Hippo pathways leads to their dedifferentiation. We used RNAi to knockdown Hth in the posterior of *Rbf GMR>yki^CA^* and examined the expression of photoreceptor markers Sens and Elav by immunofluorescence. As shown above, *Rbf GMR>yki^CA^*eye discs displayed a progressive loss of Elav expression in the posterior compartment, accompanied by the presence of ommatidia which lacked a Senseless positive R8 founder cell (Figure 4A). On average, a Sens expressing cell was missing in a quarter of ommatidia (Figure 4B). Strikingly, this phenotype was largely rescued by Hth knockdown as very few Elav positive clusters lacked a Sens positive cell (Figure 4A-B). Notably, Hth knockdown did not affect the expansion of interommatidial cells, another hallmark of Yki driven inappropriate proliferation, as the distance between ommatidia remained significantly increased in *Rbf GMR>yki^CA^ hth^RNAi^*. Additionally, there were several cases of ommatidia with duplicated R8 cells (circled in white, Figure 4A) indicating that the ability of Yki to override cell cycle exit in terminally differentiated *Rbf* mutant cells is unaffected by Hth knockdown.

**Figure 4.**
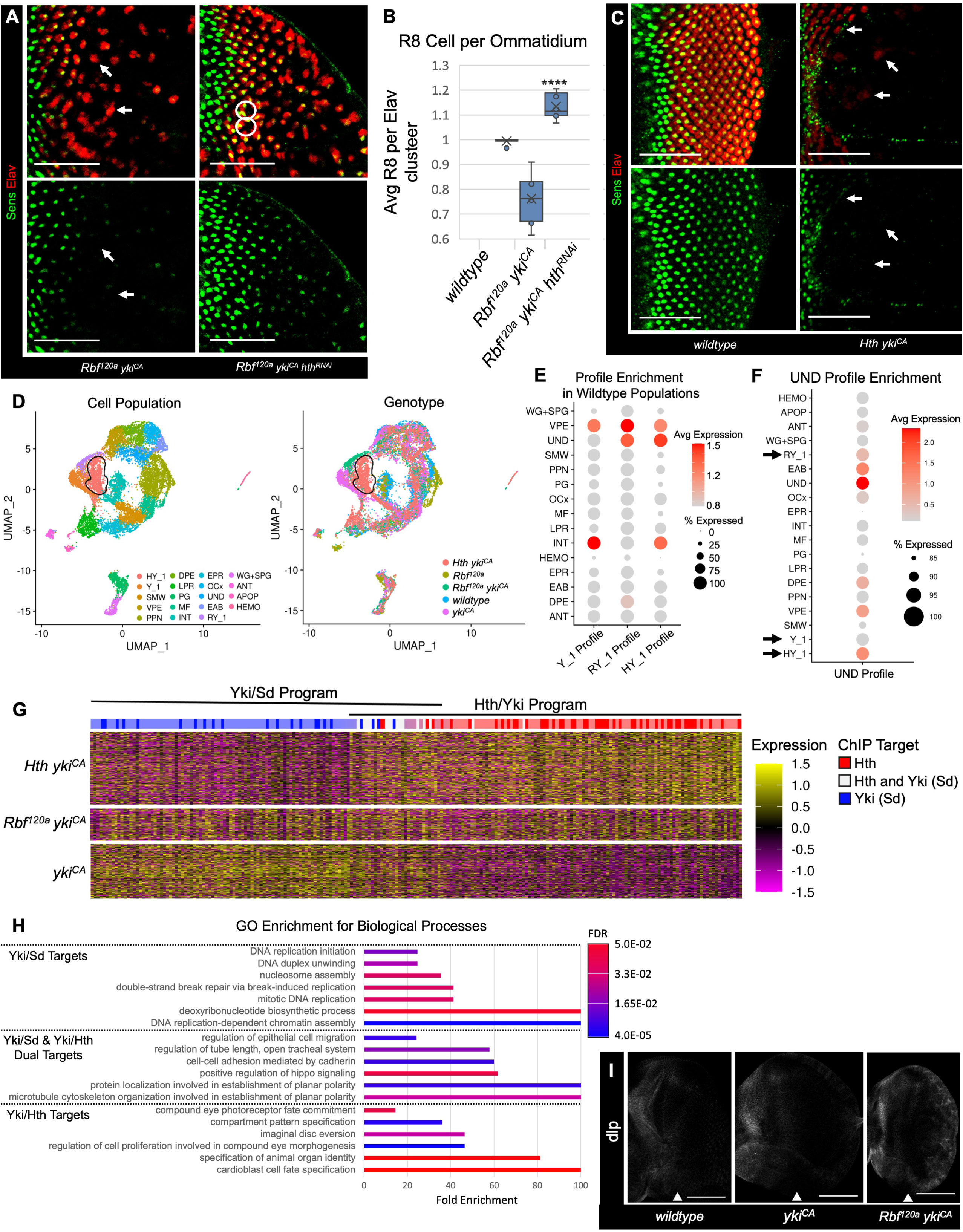
Hth is necessary and sufficient for dedifferentiation. (A) Confocal images of third instar larval eye discs from *Rbf^120a^/Y; GMR-Gal4/+; uas-yki^CA^ /+* and *Rbf^120a^/Y; GMR-Gal4/uas-hth^RNAi^; uas-yki^CA^ /+* animals stained for Elav (red) and Senseless (green). Knockdown of Hth rescues photoreceptor dedifferentiation in *Rbf GMR>yki^CA^*eye discs. Images are cropped at the MF. (B) A box-and-whisker plot displaying the ratio of Sens-positive R8 cells per Elav-positive ommatidia (N≥9 eye discs per genotype) to score the extent of dedifferentiation. (C) The *w^1118^* and *+/+; GMR-Gal4/uas-hth; uas-yki^CA^/+* eye discs stained for Elav (red) and Senseless (green). Expression of *hth* and *yki^CA^* results in photoreceptor dedifferentiation scored by the presence of Sens-negative ommatidia (white arrows). (D) An annotated UMAP of cells by population (left) and genotype-of-origin (right) in *w^1118^*, *Rbf^120a^*mutant, *GMR>yki^CA^*, *Rbf^120a^ GMR>yki^CA^*, and *GMR>hth yki^CA^* samples. A *GMR>hth yki^CA^* specific population, designated HY_1, is circled in black. (E) Dot Plot of the Y_1, RY_1, and HY_1 transcription profiles on wildtype cell populations. *GMR>yki^CA^* cells upregulate genes associated with the VPE, while *Rbf^120a^ GMR>yki^CA^*and *GMR>hth yki^CA^* cells express an UND-like signature. (F) Dot Plot of the UND transcriptional profile on all celltypes. RY_1 and HY_1 are enriched for expression of UND genes, while Y_1 is not. (G) A Heatmap of differentially expressed genes in *GMR>yki^CA^* and *GMR>hth yki^CA^* posterior cells. Genes upregulated in *GMR>yki^CA^* cells were enriched for Sd binding (dark blue lines) while *Rbf^120a^ GMR>yki^CA^*and *GMR>hth yki^CA^* were enriched for Hth target genes (red lines). Both Sd and Hth activate a subset of shared transcripts (white lines). (H) Gene ontology for biological processes (GOBP) analysis on the Yki/Sd and Yki/Hth transcriptional programs. Yki/Sd upregulate genes associated with proliferation while Yki/Hth promote expression of developmental genes. (J) Third instar larval eye discs from *w^1118^, GMR>yki^CA^,* and *Rbf^120a^GMR>yki^CA^* animals immunostained for dlp (white). *Rbf^120a^ yki^CA^* tissues aberrantly express anterior gene and Hth target dlp. Scale bars are 50 μm in all images.

To determine the impact of Hth knockdown on cell proliferation, we immunostained *GMR>yki^CA^ hth^RNAi^* and *Rbf GMR>yki^CA^ hth^RNAi^* eye discs using Elav (red) and anti-PH3 (white) antibodies to visualize photoreceptors and mitotic cells respectively (Figure S3A). Hth knockdown significantly affected the number of PH3 positive cells in neither *GMR>yki^CA^ hth^RNAi^* nor *Rbf GMR>yki^CA^ hth^RNAi^* eye discs. We then quantified the impact of Hth knockdown by analyzing the proportion of posterior PH3 positive cells in wildtype, *GMR>yki^CA^*, *GMR>yki^CA^ hth^RNAi^, Rbf GMR>yki^CA^,* and *Rbf GMR>yki^CA^ hth^RNAi^* eye discs (Figure S3B). *GMR>yki^CA^* eyes had a fourfold increase in posterior PH3 positive cells in comparison to wildtype control. There was no significant reduction in the number of mitotic cells between *GMR>yki^CA^ hth^RNAi^*and *GMR>yki^CA^* (Figure S3B). *Rbf GMR>yki^CA^* eye discs displayed over five times as many PH3 positive cells than the wildtype, and Hth knockdown in these eye discs partially reduced cell proliferation (Figure S3B). Thus, inappropriate *hth* expression in *Rbf GMR>yki^CA^* eye discs increases ectopic proliferation, but even in the absence of Hth cell proliferation remains high and is presumably driven by Yki/Sd.

To further explore the role of Hth in photoreceptor dedifferentiation, we performed a reciprocal experiment and overexpressed *hth* and *yki^CA^* with the *GMR-Gal4* driver in *Rbf* wildtype eye discs. As shown in Figure 4B, although the onset of Sens and Elav expression was not affected, the expression of both markers was acutely lost several columns posterior. Thus, Hth is necessary and sufficient to trigger photoreceptor dedifferentiation by Yki. As overexpression of *hth* can induce dedifferentiation in the presence of functional *Rbf*, this suggests that *hth* is downstream of *Rbf*.

Sd and Hth are both Yki binding partners, but their expression in wildtype tissues is different: Hth is localized to the anterior compartment, while Sd is highly expressed in the posterior compartment and at lower level in the eye anterior^32,37^. In the anterior, Hth in conjunction with Yki, inhibits differentiation and promotes the survival and proliferation of progenitor cells^34,35^. At the onset of differentiation, Yki is phosphorylated by Hippo pathway kinases leading to dramatically lower nuclear pool of Yki which, presumably, reduces Yki-dependent transcription^39^. However, ectopically active Yki in posterior cells binds to Sd and prevents cell cycle exit of interommatidial cells^36,40^. The results described above suggest that inappropriate expression of Hth in the posterior makes Hth available for Yki to execute the Yki/Hth transcriptional program in differentiating photoreceptors. To characterize how Yki/Hth affect transcription in the eye posterior, we performed scRNA-seq on *GMR>hth yki^CA^* eye discs and recovered transcriptomes of 4,004 cells (Table S7). We then performed UMAP analysis on a combined dataset of wildtype, *Rbf* mutant, *GMR>yki^CA^*, *Rbf GMR>yki^CA^* and *GMR>hth yki^CA^* cells. This revealed a population of cells unique to *GMR>hth yki^CA^*, designated HY_1 (black circle in Figure 4D). HY_1 strongly resembles RY_1, sharing over 30% of the same top markers. Further, Dot Plot analysis of HY_1 revealed similar upregulation in *bantam* and the NHR *Blimp-1* (Figure S3A). To better understand the transcriptional outcome of *hth* and *yki^CA^* overexpression, we performed differential expression analysis to generate a transcriptional profile which is characteristic of HY_1 cells (Figure S3B, Table S8). Next, we asked which wildtype cell populations have transcriptional programs most similar to that of HY_1. As shown in Figure 4E, HY_1 activates a transcriptional program that most closely resembles undifferentiated eye progenitors UND and, thus, parallels the results observed in *Rbf GMR>yki^CA^* tissue (in which *hth* is inappropriately expressed).

In a reciprocal approach, we projected the UND expression profile (Table S9) on all annotated cell types including Y_1, RY_1 and HY_1 (Figure 4F). As expected, the UND signature was readily detected in wildtype cell populations of the eye disc that have a high level of *hth* such as the EAB, VPE, and DPE. Notably, the dedifferentiating RY_1 and HY_1 clusters activated the UND signature, Y_1 did not (black arrows in Figure 4F). This further supports the idea that Yki/Sd activity alone is insufficient to induce dedifferentiation and only Yki/Hth can activate the transcriptional signature driving transformation of differentiated posterior cells into a progenitor-like cell type.

To fully characterize the transcripts activated by Yki/Sd and Yki/Hth, we then separated posterior compartment populations by genotype and performed differential expression analysis on *GMR>yki^CA^*and *GMR>hth yki^CA^* cells. This analysis produced a list of several hundred genes that were specifically upregulated in either sample. We then used this gene list to generate a Heatmap of gene expression in *GMR>yki^CA^*, *Rbf GMR>yki^CA^*, and *GMR>hth yki^CA^* cells (Figure 4G). This produced two generally distinct transcriptional programs: Yki/Sd (blue bar in Figure 4G) and Yki/Hth (red bar in Figure 4G), with a small subset of genes being activated by both Yki/Sd and Yki/Hth (Table S10). Reassuringly, the Yki/Hth transcriptional program was activated in both *Rbf GMR>yki^CA^*and *GMR>hth* yki^CA^, genotypes in which dedifferentiation was observed, but not in *GMR>yki^CA^*. We then performed motif enrichment on these expression signatures to identify transcription factors which were primarily active in each genotype^41,42^. As expected, cells from the *GMR>yki^CA^* sample were exclusively enriched for Sd target genes (dark blue bars in Figure 4G), while *GMR>hth yki^CA^* was highly enriched for Hth targets (dark red bars in Figure 4G). Interestingly, a number of the genes which were activated in both the Yki/Sd and Yki/Hth programs contained both Sd and Hth binding sites (white bars in Figure 4G), including known Yki targets such as *ex*, *Diap1*, and *kibra*^39^ and the Hippo pathway proteins *ft* and *wts*. Gene ontology of biological processes (GOBP) analysis revealed that these shared transcripts are primarily associated with cell polarity and cytoskeletal organization (Figure 4H), suggesting that regardless of binding partner Yki maintains regulation of cell morphology. Further, gene ontology for Yki/Sd specific transcripts solely consisted of GO terms for DNA replication and cell cycle (Figure 4H), reflecting that Yki/Sd drives proliferation. Uniquely, Yki/Hth program was enriched for genes with GO terms regulated to cell differentiation and fate specification, as well as proliferation (Figure 4H). This program includes upregulation in NHRs, Egfr, and several anterior-specific genes (Figure S3C).

To confirm activation of the Hth/Yki transcriptional program in *Rbf GMR>yki^CA^*, we performed immunostainings for the anterior marker and Hth target^43,44^ dally-like protein (dlp) (Figure S3F). Our single cell data show that dlp is ectopically activated in both RY_1 and HY_1. As expected, dlp was localized to the anterior compartment *wildtype* and *GMR>yki^CA,^* and was absent in the posterior (Figure 4I). Strikingly, dlp was inappropriately expressed in the posterior compartment of *Rbf GMR>yki^CA^*, further confirming a reactivation of the Hth/Yki transcriptional program in in differentiated posterior cells.

Therefore, we concluded that ectopic Hth/Yki activity triggers dedifferentiation of photoreceptors with inactivated RB and Hippo pathways. Genetic experiments demonstrate that Hth/Yki are necessary and sufficient for photoreceptor dedifferentiation, while single cell profiling confirms that reactivation of Hth/Yki in posterior cells reactivates induces the transcriptional program like that of undifferentiated eye progenitor cells.

### Rbf physically interacts with Yki and limits its ability to activate *hth* in transcriptional reporter assays

Ectopic expression of *hth* in *Rbf GMR>yki^CA^*eye discs is dependent on Sd (Figure 3E), a DNA-binding protein that recruits Yki to its target genes^32,38^, raising the possibility that Yki directly regulates *hth* expression. Therefore, we examined the occupancy of Yki and Sd at the *hth* locus using publicly available genome-wide binding data from the larval wing disc, eye disc, and embryos^21,45,46^. Indeed, both Yki and Sd were enriched at multiple regions within the *hth* gene, with closely matching peaks (Figure 5A). Another important factor in Yki dependent transcription is the GAGA transcription factor GAGA (GAF), which physically interacts with Yki and contributes to Yki activation^20,21^. Interestingly, GAF showed significant colocalization with Yki/Sd peaks throughout the *hth* gene. To explore the role of these proteins in *hth* regulation, we generated a series of luciferase reporter constructs containing regions of the *hth* gene that showed enrichment for Yki, Sd, and GAF occupancy. The luciferase reporters were then transfected into Drosophila S2R+ cells and their response to co-transfected Yki/Sd, GAF and Rbf was determined. The reporter containing fragment within a black rectangular in Figure 5A was modestly activated by Yki/Sd, while Rbf limited this activation (Figure 5B). Although GAF was previously shown to potentiate Yki activation of its targets, unexpectedly, GAF co-transfection blocked Yki/Sd activation of the reporter. The repression of the reporter was further potentiated by Rbf co-expression.

**Figure 5.**
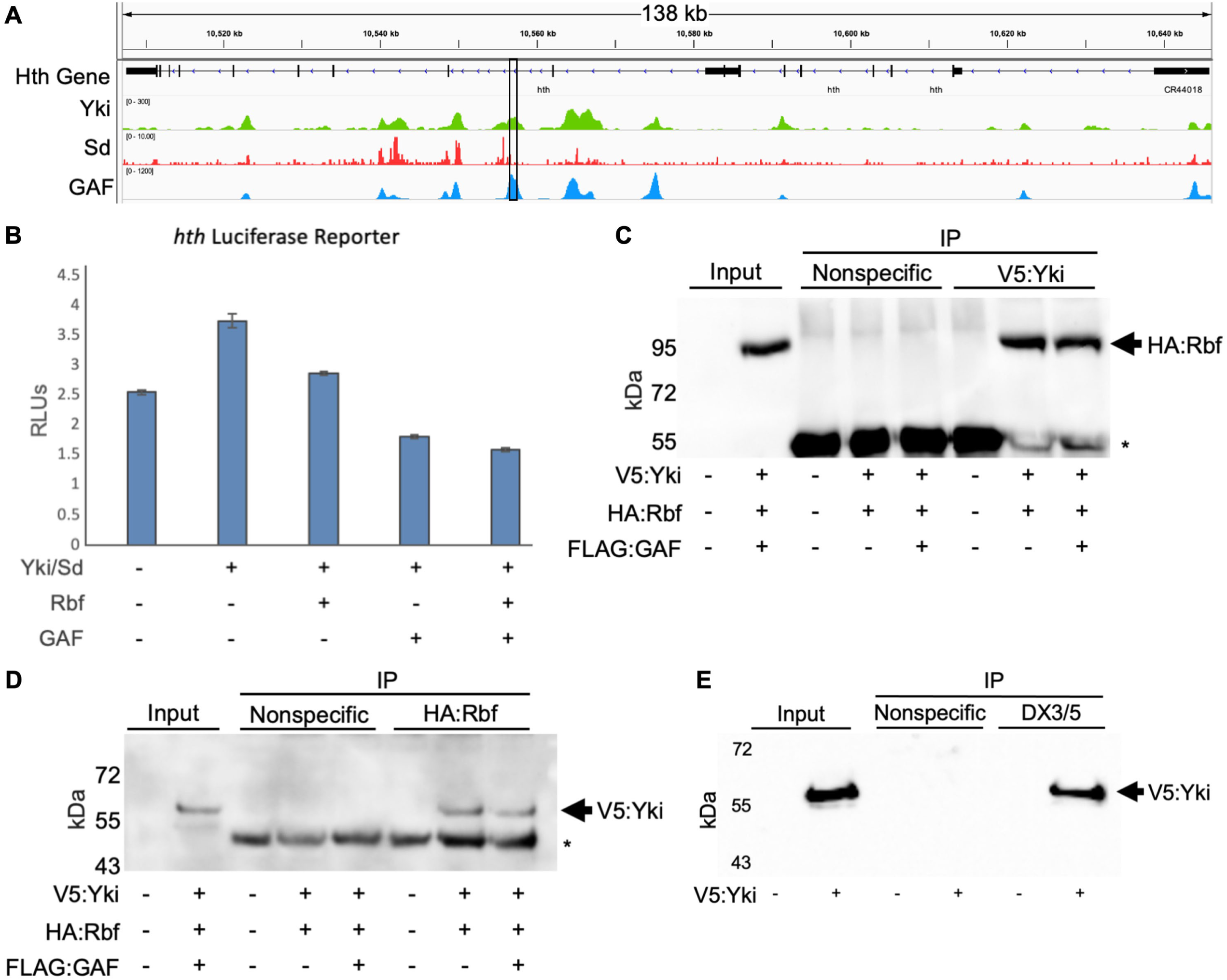
Rbf and Yki physically interact to repress the expression of the *hth* reporter. (A) ChIP-Seq data showing Yki, Sd, and GAF occupancy at the *hth* gene. The position of the fragment used to make a *hth-*Luc reporter is outlined in black. (B) S2R+ cells were transfected with a *hth* luciferase reporter and to in S2R+ cells expressing combinations of yki, Sd, GAF, and Rbf. Yki/Sd upregulate *Hth* which is suppressed by the addition of Rbf and GAF. (C) S2R+ cells were transfected with V5:Yki, HA:Rbf, and FLAG:GAF. Lysates were immunoprecipitated with mouse anti-CD133 as a nonspecific negative control and anti-V5 to detect V5:Yki. Western blot was probed with with anti-HA to visualize HA:Rbf. Asterisks denote antibody heavy chains. (D) S2R+ cells were transfected with V5:Yki, HA:Rbf, and FLAG:GAF. Lysates were immunoprecipitated with mouse anti-CD133 as a nonspecific negative control and anti-HA to detect HA:Rbf. Western blot was probed with anti-V5 to visualize V5:Yki. Asterisks denote antibody heavy chains. (E) Endogenous Rbf physically interacts with V5-Yki. S2R+ cells were transfected with V5:Yki and lysates were immunoprecipitated with mouse anti-CD133 as a nonspecific negative control and DX3/5 antibodies to immunoprecipitate endogenous Rbf. Western blot was probed with anti-V5 to visualize V5:Yki.

Given that Rbf limited Yki/Sd activation of the reporter, we tested whether Rbf and Yki physically interact. V5-Yki and HA-Rbf were transiently transfected in S2R+ cells, the complexes were immunoprecipitated with V5 antibody, separated by SDS-PAGE, and western blots were probed with HA antibody to detect HA-Rbf. As shown in Figure 5C, HA-Rbf was significantly enriched in the V5-Yki immunoprecipitates while no HA-Rbf was immunoprecipitated in the negative control. Co-transfection of GAF did not affect the interaction between HA-Rbf and V5-Yki. Next, we performed a reciprocal experiment and confirmed the presence of V5-Yki in HA-Rbf immunoprecipitates (Figure 5D). Finally, cells were transfected with V5-Yki and endogenous Rbf was immunoprecipitated from the cell extracts. The presence of V5-Yki in Rbf immunoprecipitates was detected by western blotting with V5 antibody (Figure 5E). Thus, we concluded that Yki associates with Rbf.

### Knockdown of GAF leads to ectopic *hth* expression in *GMR>yki^CA^* eye disc

GAF has been shown to promote Yki/Sd gene expression in the context of cell proliferation^20,21^. Therefore, finding that GAF limits activation of the *hth-Luc* reporter by Yki/Sd was unexpected. To explore the effect of GAF on *hth* expression in endogenous settings, we used RNAi to deplete GAF (*Trl*) in the posterior compartment of the *GMR>yki^CA^* eye disc and determined Hth expression by immunofluorescence (Figure 6A). The *UAS*-*Trl^RNAi^* transgene has been previously validated to efficiently knockdown GAF protein in vivo and studied in the context of Yki dependent transcription^20^. Strikingly, Hth was inappropriately expressed posterior to the MF in *GMR>Trl^RNAi^ yki^CA^* at a level similar to *Rbf GMR> yki^CA^* (Figure 6B). This upregulation was accompanied by progressive reduction in the number of Elav positive cells across the eye posterior, indicating that photoreceptors in *GMR>Trl^RNAi^ yki^CA^* undergo dedifferentiation. Accordingly, staining with Elav and Sens antibodies revealed the appearance of Elav positive clusters that lack a Sens positive cell (white arrows in Figure 6C), a hallmark of dedifferentiation.

**Figure 6.**
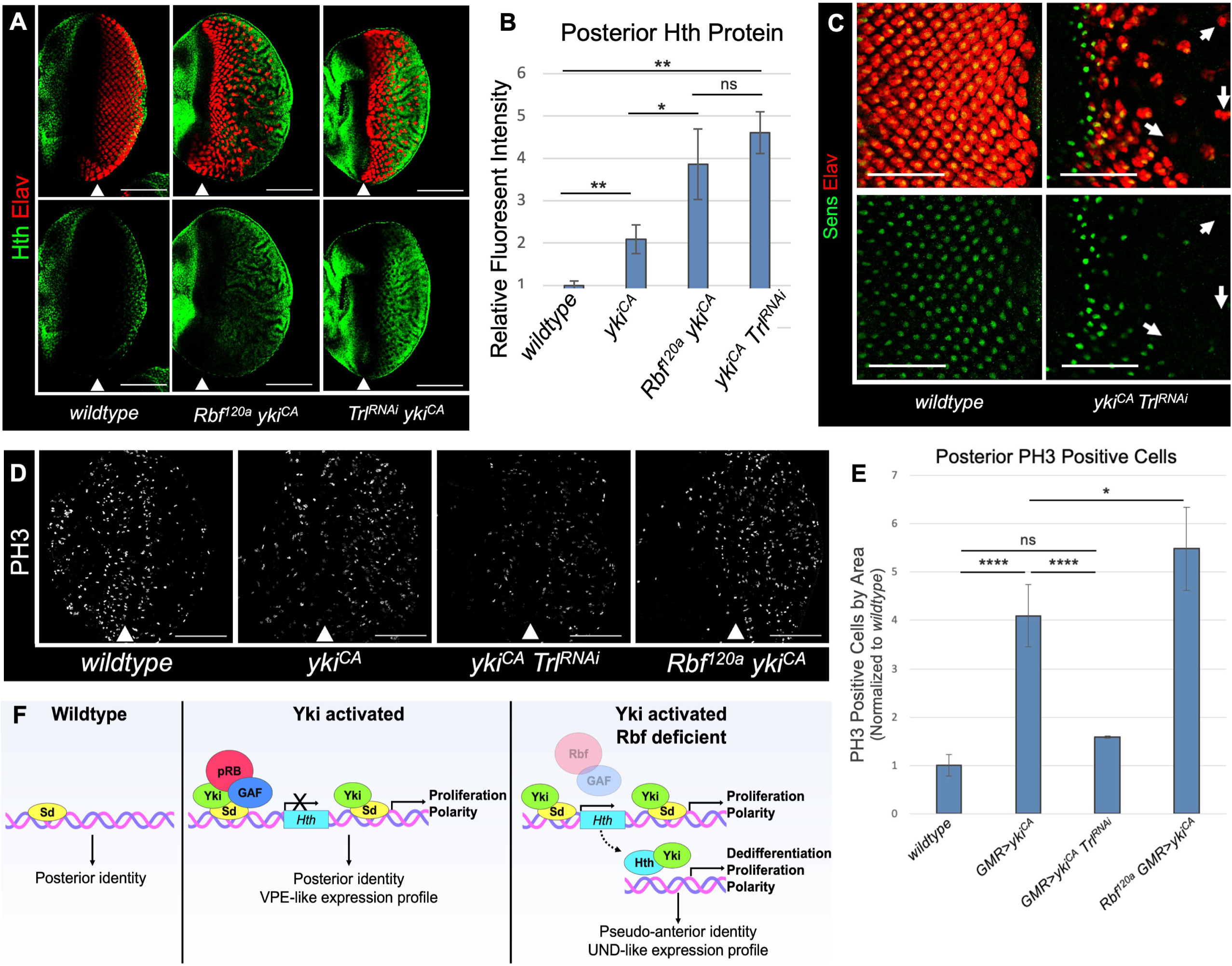
Depletion of GAF derepresses *hth* expression and triggers dedifferentiation in Yki expressing photoreceptors. (A) *Trl* knockdown derepresses Hth in the posterior of *GMR>yki^CA^*eyes. Larval eye discs from *w^1118^*, *Rbf^120a^/Y;GMR-Gal4/+;uas-yki^CA^/+*, and *w^1118^/Y;GMR-Gal4/uas-Trl^RNAi^; uas-yki^CA^ /+* animals were stained for Elav (red) and Hth (green) antibodies. (B) A bar graph displaying Hth protein levels in the posterior compartment normalized to the anterior compartment of each tissue (N≥6 per genotype). (C) *GMR>Trl^RNAi^ yki^CA^* photoreceptors fail to maintain neuronal specification as visualized by the occurrence of Elav-positive, Sens-negative ommatidial clusters (white arrows). (D) *Trl* knockdown blocks *yki^CA^* driven inappropriate proliferation. Eye discs from *w^1118^*, *w^1118^/Y;GMR-Gal4/+;uas-yki^CA^ /+*, *w^1118^/Y;GMR-Gal4/uas-Trl^RNAi^;uas-yki^CA^/+*, and *Rbf^120a^/Y;GMR-Gal4/+;uas-yki^CA^/+* animals were stained for Elav (red) and PH3 (purple). (E) A bar graph displaying the number of mitotic, PH3-positive cells in the posterior compartment of each genotype. (F) A model for RB and Hippo pathways cooperation in maintenance of the differentiation state. In Rbf proficient Hippo pathway mutant cells, Rbf blocks Yki/Sd from activating *hth* in the eye posterior. In this cells, Yki/Sd drives the expression of proliferation and polarity genes and infers a ventral peripodial like signature. In the absence of Rbf, Yki/Sd inappropriately activates *hth* in differentiating photoreceptors. The availability of Hth results in activation of the Yki/Hth transcriptional program that resembles the transcriptional signature of the eye progenitor cells (UND) and leads to photoreceptors dedifferentiation. Thus, in addition to the canonical Yki/Sd signature, RB and Hippo pathways double mutant cells activate the Yki/Hth transcriptional program. Scale bars are 100 μm in all images.

To further dissect the proliferation and dedifferentiation phenotypes, we then stained *Trl* knockdown *yki^CA^* expressing eye discs for PH3 to visualize mitotic cells. As expected, knockdown of GAF strongly reduced *yki^CA^* driven proliferation in the context of Hth expression (Figure 6D). In fact, GAF knockdown successfully limited proliferation to wildtype levels in *yki^CA^* expressing cells (Figure 6E). Thus, although GAF is necessary for activation of the Yki/Sd pro-proliferative program it acts as a repressor at the *hth* gene.

## DISCUSSION

Here, we describe a mechanism by which the RB and Hippo pathways cooperate to maintain photoreceptor differentiation, a function which is distinct from the role of Hippo in cell fate specification. In Hippo pathway deficient cells with normal RB function, ectopic Yki activates a Sd-dependent pro-proliferative program which is characteristic of peripodial membrane cells^39^. This results in gross expansion of interommatidial cells and may affect cell fate but does not impact photoreceptor identity once specified^22,39^. In contrast, combined inactivation of RB and Hippo pathways, in addition to activation of the Sd/Yki program, leads to inappropriate expression of the progenitor gene *hth* in differentiating photoreceptors in a Yki/Sd dependent manner. We suggest that availability of Hth now allows Yki to execute a distinct, Hth/Yki transcriptional program that no longer resembles that of peripodial cells but is instead characteristic of eye progenitor cells (Figure 6F). This program imparts previously specified photoreceptor neurons with an eye progenitor-like phenotype akin to undifferentiated cells of the anterior compartment. Thus, the RB and Hippo tumor suppressor pathways maintain photoreceptor identity by restricting Sd/Yki dependent expression of *hth* to prevent reactivation of the progenitor transcriptional program in terminally differentiating cells.

In the wildtype eye, *hth* is highly expressed in the anterior compartment^47^ where it, together with Tsh, promotes proliferation of eye progenitor cells and suppresses premature photoreceptor differentiation. Hth activity in the anterior is crucial for maintaining an undifferentiated state and its loss in these cells results in ectopic differentiation of photoreceptors. It has been shown that Hth and Tsh promote proliferation of these eye progenitor cells, at least partially, by partnering with Yki^32^. Accordingly, artificially prolonging *hth* expression in clones of eye imaginal tissue inhibits MF initiation, suppresses photoreceptor recruitment and abrogates neuronal differentiation^34,35^. Ectopic activation of hyperactive *yki^S168A^* in undifferentiated tissues has been shown to induce expression of *wg* which promotes *hth/tsh* expression to inhibit photoreceptor differentiation^48^. Further, expression of *yki* in developing eye discs under certain conditions can impair eye development and produce tissues which are completely devoid of photoreceptor differentiation or patterning^39,49^. The common theme of these studies is that ectopic Yki is able, in some contexts, to prevent the onset of photoreceptor specification by interfering with the developmental cues to differentiate. Therefore, these defects are explicitly distinct from the phenotype of *Rbf wts* double mutant cells, which initiate proper execution of the neuronal differentiation program, establish axonal projections, but fail to maintain a differentiated state^17^. Genetic tracing experiments now conclusively show that *Rbf wts* double mutant photoreceptors indeed lose their neuronal identity and are not eliminated, in agreement with our earlier marker analysis^17^. Thus, Hippo and RB pathways cooperate to retain cells in a state of terminal differentiation.

Several lines of evidence suggest that inappropriate expression of *hth* in the eye posterior is the key event in photoreceptor dedifferentiation. Ectopic expression of *hth* was observed in three genetic experimental settings that were accompanied by a progressive loss of photoreceptor identity, as evident by the appearance of ommatidia lacking an R8 founder cell. Accordingly, depletion of Hth in *Rbf* mutant photoreceptors expressing constitutively active *yki^CA^* rescues photoreceptors from dedifferentiation, while co-expression of *hth* with *yki^CA^* in *Rbf* wild type eye discs robustly induces dedifferentiation. Thus, *hth* is necessary and sufficient for dedifferentiation of photoreceptors that express *yki^CA^* and is downstream of *Rbf*. Notably, depletion of Hth rescued dedifferentiation defects in *Rbf GMR>hth^RNAi^ yki^CA^* but not ectopic cell proliferation, which is in agreement with previous findings that photoreceptor dedifferentiation is independent of ectopic proliferation^17^.

Why does inappropriate upregulation of *hth* in the presence of activated Yki result in dedifferentiation? Our single cell data strongly argue that photoreceptor dedifferentiation is specifically accompanied by activation of a distinct transcriptional program that is most similar to that of the eye progenitor cells anterior to the MF and is replete with Hth target genes. Activation of Hth target genes occurs in both *Rbf GMR>yki^CA^* and *GMR>hth yki^CA^* settings and is absent in cells expressing *yki^CA^* alone. This program is activated in addition to the classical Yki/Sd transcriptional program suggesting that availability of Hth does not prevent Yki from partnering with Sd. In line with previous work, projection of the *yki^CA^* expression profile onto wildtype cells revealed that Yki/Sd upregulate a cassette of genes typically restricted to the peripodial membrane^39,49^. Interestingly, a number of known Yki targets such as *Diap1*, *ex*, and *kibra*, and the Hippo pathway components *ft* and *wts* are shared between the Yki/Sd and Yki/Hth programs. Many of these genes are direct targets of both Sd and Hth, while gene ontology analysis for these shared targets revealed enrichment GO terms for cytoskeletal organization and cell polarity, suggesting that regardless of binding partner Yki is highly influential in regulation of cell morphology and adhesion. Notably, overexpression of Hth alone in the posterior compartment is inconsequential and requires Yki to trigger dedifferentiation. Given that Hth and Yki physically interact and were shown to regulate Hippo pathway targets such as *bantam*^32^, this data collectively suggests that the dedifferentiation program is driven by Yki/Hth.

Genome wide chromatin binding experiments revealed that *hth* is a direct target of Yki/Sd^21,46^. Accordingly, Yki/Sd activates the *hth* luciferase reporter and this activation is blocked by Rbf. Given that Rbf is a known transcriptional repressor that blocks E2F activation through direct binding, our observation that Rbf similarly interacts with Yki provides a simple explanation for why Yki does not activate *hth* in Rbf proficient cells. The finding that GAF similarly inhibits Yki/Sd expression of *hth* was especially surprising. Previous studies have shown that GAF is necessary for full activation of the Yki/Sd proliferation program^20,21^. However, in the context of *hth* regulation, GAF appears to negatively regulate its expression both *in vivo* and in reporter assays in cultured cells. Thus, GAF appears to simultaneously promote Yki/Sd driven proliferation and protect these proliferating cells from dedifferentiation through inhibition of *hth*.

The idea that Rbf represses the expression of genes determining the progenitor state, such as *hth*, is consistent with the role of mammalian pRB. pRB has been previously shown to repress transcription of key pluripotency genes Sox2 and Oct4 in somatic cells^6^. Further, RB protects prostate cancer cells from lineage transformation through inhibition of Sox2 and chromatin modifying protein Ezh2^50^. Thus, the cooperation of Hippo and RB pathways in maintaining a state of differentiation converges on their regulation of *hth*, where the Hippo kinase cascade and Rbf limit Yki activity.

## STAR Methods

### Fly Stocks and Maintenance

Stocks and crosses were kept at 25C in vials containing standard Bloomington formulation media.

*w^1118^* was used as a wildtype stock for all single-cell RNA-Sequencing and genetic experiments.

*Rbf^120a^/FM7, GFP* (gift from Nick Dyson)

yw, ey-Flp/FM7, GFP; uas-mFlp5, Actin5c, mFRT71C, stop, mFRT71C, lacZ^nls^/CyO, GFP; elav-GAL4, FRT82B, GFP/TM6B

Rbf^120a^, ey-Flp/FM7, GFP; uas-mFlp5, Actin5c, mFRT71C, stop, mFRT71C, lacZ^nls^/CyO, GFP; elav-GAL4, FRT82B, GFP/TM6B

*Rbf^120a^, ey-Flp/FM7, GFP;;FRT82B GFP/TM6B w*; FRT82B, wts^x1^/TM6B* (BDSC 9146)

*w^1118^; GMR-GAL4/CyO* (BDSC 9146)

*w*;; uas-yki ^S111A, S168A, S250A,V5^*(BDSC 28817)

*uas-Sd^RNAi^* (VDRC ID 1014978)

*uas-Hth^RNAi^* (VDRC 100630)

*uas-Rbf^RNAi^* (VDRC ID 10696)

*uas-Trl^RNAi^* (BDSC 67265)

*uas-Trl^RNAi^* (VDRC ID 106433)

*uas-hth* (gift from Richard Mann)

Final Stocks:

yw, ey-Flp/+; uas-mFlp5, Actin5c, mFRT71C, stop, mFRT71C, lacZ^nls^/+; elav-GAL4, FRT82B, GFP/FRT82B, wts^x1^

Rbf^120a^, ey-Flp/Y; uas-mFlp5, Actin5c, mFRT71C, stop, mFRT71C, lacZ^nls^/+; elav-GAL4, FRT82B, GFP/FRT82B, wts^x1^

yw, ey-Flp/+;; FRT82B, GFP/FRT82B, wts^x1^

Rbf^120a^, ey-Flp/Y;; FRT82B, GFP/FRT82B, wts^x1^

w*^1118^*/+; GMR-GAL4/+; uas-yki*^S111A, S168A, S250A,V5^*/+

*Rbf^120a^/Y; GMR-GAL4/+; uas-yki ^S111A, S168A, S250A,V5^/+*

*w^1118^/+; GMR-GAL4/uas-hth^RNAi^; uas-yki ^S111A, S168A, S250A,V5^/+*

*Rbf^120a^/Y; GMR-GAL4/uas-hth^RNAi^; uas-yki ^S111A, S168A, S250A,V5^/+*

*w^1118^/+ or Y; GMR-GAL4/uas-Trl^RNAi^; uas-yki ^S111A, S168A, S250A,V5^/+*

### Immunofluorescence

Eye imaginal discs were dissected from wandering third instar larvae, or pupa when specified, and dissected in ice-cold 1X PBS. Samples were then fixed in 4% formaldehyde (Polysciences 18814) diluted in room temperature 1X PBS for 30 minutes and subsequently permeabilized in 0.3% PBS-Triton X-100 twice for 10 minutes. Blocking was performed for one hour at room temperature in 10% normal donkey serum (Jackson ImmunoResearch 017-000-121) with 0.01% Triton X-100. Samples were then incubated in primary antibody diluted in 1X PBS with 10% NDS overnight at 4C on a nutator. The following day, samples were washed three times for 5 minutes in 0.1% PBS-Triton X-100 before incubating in either Cy3 or Cy5-conjugated secondary antibodies (Jackson ImmunoResearch 1:200) for 1 hour at room temperature including 4′,6-diamidino-2-phenylindole (DAPI). Finally, samples were washed four times with 0.1% PBS-Triton X-100 for five minutes before mounting on glass slides in FluorSave reagent (MilliporeSigma 345789). Primary antibodies and concentrations include: anti-β-Gal (DSHB 40-1a, 1:200), anti-Elav (DSHB 7EA10, 1:200), anti-Senseless (from H. Bellen, 1:2000), anti-Hth (from Richard Mann, 1:2000), anti-PH3 (Millipore, 1:500), and anti-dlp (DSHB 13G, 1:50).

### Microscopy

A Zeiss Axio Observer Z1 LSM 700 confocal microscope was used to capture fluorescent images at x20/0.8 and x100/1.45 objectives. All images were taken using 1 AU pinhole and gain settings were consistent for all images within an experiment. Resulting images were processed using ImageJ Software (1.48v, NIH, USA) unless otherwise specified. Image processing included adjustments to brightness and contrast which were consistent for all images in each experiment. Images shown are representative.

### Tissue Dissection for scRNA-Seq

Third instar larvae were harvested upon wandering and eye imaginal discs were dissected in cold 1X PBS. Eye discs were then dissected from the brain and antenna by cutting the optic stalk and eye-antennal-border using a microblade. Discs were then deposited in ice-cold 1X Rinaldini solution. Collagenase (Sigma #C9891) was added to achieve a final concentration of 2.5 mg/mL and Trypsin (Sigma #59418C) to 1X dilution. Tubes were then attached horizontally to a shaker and agitated at 225 rpm for 10 minutes at 37C. Samples were then subjected to approximately 30 pulses of gentle mechanical digestion using a P200 pipette. The single-cell suspension was then washed twice in 0.04% BSA-PBS and resuspended in 30 uL of 0.04% BSA-PBS. The concentration of cells and their viability was assessed using a hemocytometer and Trypan blue staining.

### Hth Protein Quantification

Using ImageJ software, confocal images of eye imaginal discs were cropped to remove antennal discs before defining the anterior and posterior compartments. The DAPI channel was then used to outline the border of the tissue while Elav signal was used to define the posterior compartment. Hth protein defined the anterior compartment. In mosaic tissues, GFP signal was used to further separate *wts* wildtype and mutant tissue. The average fluorescent intensity of the Hth channel was measured in the anterior and posterior compartments from 6-10 tissues across multiple experiments using the ImageJ (1.48v, NIH, USA) measure function (Analyze- >Measure). The average Hth fluorescent intensity of each posterior compartment was then normalized to the anterior compartment of the same sample to account for differences across experiments. These ratios were then averaged to reflect the overall abundance of Hth in the posterior compartment in each genotype. Hth abundance ratios were then pooled and analyzed by Student’s T-Test to assess significance (ns=p>0.05, *= p≤0.05, **= p≤0.01, ***= p≤0.001, ****p≤0.0001). Data is presented as a bar graph with error bars representing the standard deviation of samples across each genotype.

### Photoreceptor Differentiation Quantification

Using ZEN Blue 2.6 software (ZEISS), Elav positive cells were used to define ommatidia into zones-of-influence (ZOI). Gaussian smoothing was then applied to better capture all photoreceptors in a single ommatidium and binary erosion was selected to separate adjacent ommatidia. R8 cells were then identified within each ommatidium by measuring the presence of Senseless signal within its corresponding ZOI. Background fluorescence was omitted by restricting the minimum area for senseless-positive cells to 25 pixel units. Thresholds were adjusted for each image to account for variations in fluorescent intensity across samples and experiments. Data was condensed to reveal the number of ommatidia and R8 cells identified per tissue. Between 9 and 10 samples were processed for each genotype and Student’s T-Test was performed to assess significance. Results are displayed as a box-and-whisker plot.

### PH3 Quantification

Using ImageJ software, confocal images of eye tissues were cropped to the posterior compartment using Elav immunostaining. The pixel area of each posterior compartment was measured using the Measure function (Analyze->Measure). Fluorescent intensity thresholds for the PH3 channel were set to 50-255 (Image->Adjust->Threshold) before running Analyze Particles (Analyze->Analyze Particles) with size thresholds set from 5-infinity pixel units to count the number of PH3 foci. The number of PH3 positive foci was then normalized to the sampled area of the posterior compartment and averaged for each genotype (n=5-10 samples per genotype). The significance of the ratio of PH3 foci per area was assessed by Student’s T-Test (ns=p>0.05, *= p≤0.05, **= p≤0.01, ***= p≤0.001, ****p≤0.0001). Data is presented as a bar graph with error bars representing the standard deviation of samples across each genotype.

### Identification of Protein Binding Sites

ChIP-seq data from Hth^43^ (SRX111801), Yki and GAF^21,51^ (GSE38594), Sd^52^ (GSE54601), and Rbf and Dp^3^ (GSE102043) were loaded into IGV^53^ (version 2.94) for visualization, analysis, and peak identification.

### Construction of Luciferase Vectors

*Hth* gene regions containing Yki, Sd, and GAF binding peaks were PCR amplified from *w^1118^* genomic DNA. PCR products were then cloned into the XhoI site of a pGL3 luciferase reporter vector (Promega) including an hsp70 minimal promoter^18^ using NEBuilder Hifi DNA Assembly (NEB). Appropriate insert sequences were confirmed by Sanger sequencing. Primers and respective gene products cloned into luciferase reporters are listed in Table S11.

### Luciferase Assay

S2R+ cells were plated onto 24-well plates at 10*10^3^ cells per well in 10% FBS-Schneider’s Drosophila Medium (Sigma S0146-500ML) and allowed to adhere for 1 hour at 25C. For each well, 10 μg of luciferase reporter vector and 1 μg of renilla reporter were transfected with a combination of empty vector, 200 ng Yki, 200 ng Sd, 200 ng Rbf, or 200 ng GAF, for a total of 811 ng per well. Transfections were performed using 50 μL serum-free media containing 2 μL X-Tremegene HP DNA Transfection Reagent (Sigma 6366244001) and incubated for 48 hrs at 25C. After transfecting, cells were washed in cold PBS and lysed in 100 μL 1X Passive Lysis Buffer (Promega E1941). In duplicate, 25 uL of each lysate was loaded on a white, flat-bottom 96-well plate and subjected to the Dual-Luciferase® Reporter Assay System (Promega E1910). Luciferase assays were performed at least 3 times.

### Immunoprecipitation-Western Blot

Drosophila S2R+ cells were plated into 6-well plates at 2*10^8^ per well in 10% FBS-Schneider’s Drosophila Medium (Sigma S0146-500ML) and allowed to adhere for 1 hour at 25C. For each well, 3 μg of pIEX-7 Ek/LIC vector (Novagen) encoding either empty vector or epitope tagged proteins were incubated in 100 μL serum-free media containing 2 μL X-Tremegene HP DNA Transfection Reagent (Sigma 6366244001) for 30 minutes at room temperature. After adhering, cells were transfected with plasmid and incubated for 48 hours at 25C. Post-transfection, cells were harvested and snap-freezed in Dynlacht buffer containing 1X protease inhibitor (Sigma 11836170001) before incubating overnight in primary antibody at 4C. Mouse anti-CD133 (DSHB HB#7, 1:100) was used as a negative control, mouse anti-V5 for V5-tagged Yki (Fischer R960-25, 1:100), mouse anti-HA for HA-tagged Rbf (Fischer 26183, 1:100), and mouse anti-Rbf (DX3 and DX5, 1:10) for endogenous Rbf.

The following day, lysates were incubated in 20 μg Dynabeads Protein G and A 1:1 (Fischer 10003D and 10001D). Dynabeads were then washed four times for 10 minutes in cold Dynlacht buffer before eluting immunoprecipitates into Laemmli buffer and separating by SDS-PAGE. Resulting Western Blots were blocked for 30 minutes at room temperature in 2.5% non-fat dry milk TBS with 0.1% Tween-20 before incubating in mouse anti-V5 (Fischer R960-25, 1:2000) or mouse anti-HA (Fischer 26183, 1:2000) overnight at 4C. The following day, blots were washed three times for 5 minutes with TBS-T before incubating in anti-mouse secondary conjugated to HRP (1:3000) for one hour. Samples were then washed three times for five minutes in TBS-T, incubated in SuperSignal Chemiluminescent substrate for 5 minutes (ThermoFischer 34577), and imaged on Azure imaging system. Resulting images were processed through adjustments to brightness and contrast and horizontal cropping for clarity.

### scRNA-Seq Data Preprocessing

The single cell sample Fastq files were processed using the 10X Genomics CellRanger (version 6.0.0) count function with default parameters. Reads were aligned to *Drosophila* genome BDGP6 which was extracted from Ensemble (version 90). Features, barcodes, and matrices generated by CellRanger were then processed into Seurat objects using the Read10X and CreateSeuratObjects functions. Data was deposited to the NCBI Gene Expression Omnibus database (GEO, https://www.ncbi.nlm.nih.gov/geo/) under accession number GSE217380.

### scRNA-Seq Data Analysis

Datasets were merged using the Seurat R package^54–56^ and low viability cells were removed on the basis of low RNA content and enrichment for mitochondrial, heat shock, and ribosomal genes. Remaining cells were processed using default settings. UMAPs were created using 45 PCs before differential expression analysis with FindAllMarkers using the Wilcoxon Rank Sum. LogFC thresholds and min.pct, the minimum fraction of cells expressing a given gene, were both set to 0.25. Populations were then annotated based on top marker expression as described in Ariss 2018^29^. Novel populations were characterized by their genotype of origin. Specific transcriptional profiles were generated from differential expression analysis of the population-of-interest versus all other cells, combined with previously determined top markers. For example, the posterior compartment transcriptional profile was generated using differential expression of all posterior cells versus all anterior cells. This list was then supplemented with top marker genes for known posterior cell types which were not captured during pseudobulk differential expression analysis. Enrichment for expression of these transcriptional profiles was determined using AddModuleScore and resulting scores were visualized using the FeaturePlot, DotPlot, and DoHeatmap functions.

### Cell Trajectory Analysis

Single-cell trajectory analysis was performed using Monocle 3^31^. Seurat objects and UMAPs were converted into Monocle compatible celldataset objects and UMAP pseudotime was calculated using the branch point within the EAB cluster, the least differentiated population, as the root node.

### Gene Ontology Analysis

PANTHER Overrepresentation Test for GO biological processes in *Drosophila melanogaster* was performed using Gene Ontology Resource^57–59^ (http://geneontology.org) with default settings and FDR P < 0.05. Fold enrichment and FDR values were then input to Microsoft Excel for visualization.

### Motif Enrichment Analysis

Motif enrichment for transcription factor binding to differentially expressed genes was performed with i-cisTarget^41,42^ (database version 6.0) using the dm6 *Drosophila* genome. Genes were analyzed against a Transcription Factor Binding Site ChIP database containing data from 436 ChIP-seq experiments. Data was submitted with default settings: a 0.4 minimum fraction of overlap, NES threshold of 3.0, ROC threshold of 0.01, and a visualization threshold of 5000. Candidate target genes for relevant ChIP experiments, including Sd ChIP-seq on eye-antennal discs (SRX457598) and hth ChIP-seq on 8-16h embryos (SRX111801), were then extracted.

## Data Availability

scRNA-seq data generated for this publication is deposited in the GEO database (https://www.ncbi.nlm.nih.gov/geo/) under accession number GSE217380.

## Acknowledgements

Our thanks to Paula Zappia for her discussions and advice on single-cell RNA-sequencing and data analysis and to James Kwon for his assistance in constructing the *hth* luciferase reporter. We thank Kenneth Irvine for providing the V5:Yki plasmid, Hugo Bellen for anti-Senseless antibody, Richard Mann for anti-Hth antibody and UAS-*hth* flies and the Developmental Studies Hybridoma Bank at the University of Iowa for anti-Elav, anti-β-Gal, and anti-dlp antibodies. Other stocks were obtained from the Bloomington Drosophila Stock Center (NIH P40OD018537) and Vienna Drosophila Resource Center. Finally, we are grateful to Flybase for information and resources regarding the *Drosophila* genome. This work was supported by NIH grant R35GM131707 to M.V.F.

## Author Contribution

Conceptualization, AER and MVF; Methodology, AER and BB. Investigation, AER and BB. Visualization, AER. Supervision, MVF. Writing-Original Draft, AER. Writing-Review and Editing, AER and MVF. Funding Acquisition, MVF.

## Supplemental Tables

**Table S1: *Rbf wts* scRNA-seq experiment expression matrix**

**Table S2: Anterior vs. posterior differentially expressed genes**

**Table S3: The RW_1 transcriptional profile**

**Table S4: *Rbf yki* scRNA-seq experiment expression matrix**

**Table S5: The Y_1 transcriptional profile**

**Table S6: The RY_1 transcriptional profile**

**Table S7: *Hth yki* scRNA-seq experiment expression matrix**

**Table S8: The HY_1 transcriptional profile**

**Table S9: The UND transcriptional profile**

**Table S10: *GMR>yki^CA^* vs. *GMR>hth yki^CA^* shard and differentially expressed genes**

**Table S11: Primer sequences and gene products for luciferase constructs**

## Supplemental Figure legends

**Figure S1.**
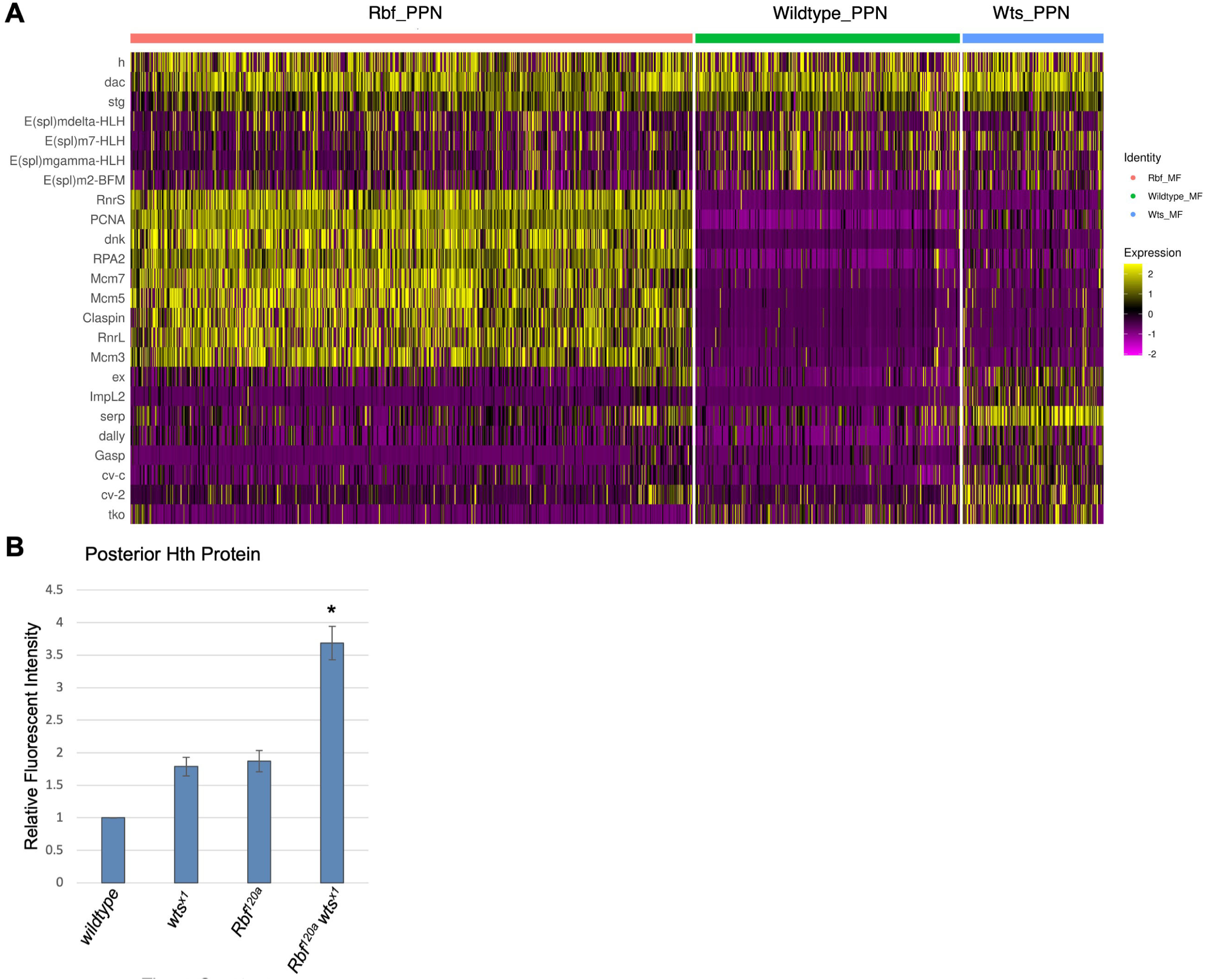
Rbf and wts single mutations impact PPN transcription, do not upregulate *hth*. (a) A Heatmap of genes differentially expressed between *w^1118^, wts,* and *Rbf^120a^ wts^x1^* PPN populations. *Rbf^120a^* mutant PPN cells upregulate proliferative genes and *wts^x1^*mutant PPN cells overexpress Yki target *ex*. Both maintain expression of PPN markers. (b) A bar graph displaying levels of Hth protein in the posterior compartment normalized to the anterior compartment of each tissue (N≥4). There is a significant increase in Hth protein in the posterior compartment of *Rbf^120a^ wts^x1^*double mutant tissue.

**Figure S2.**
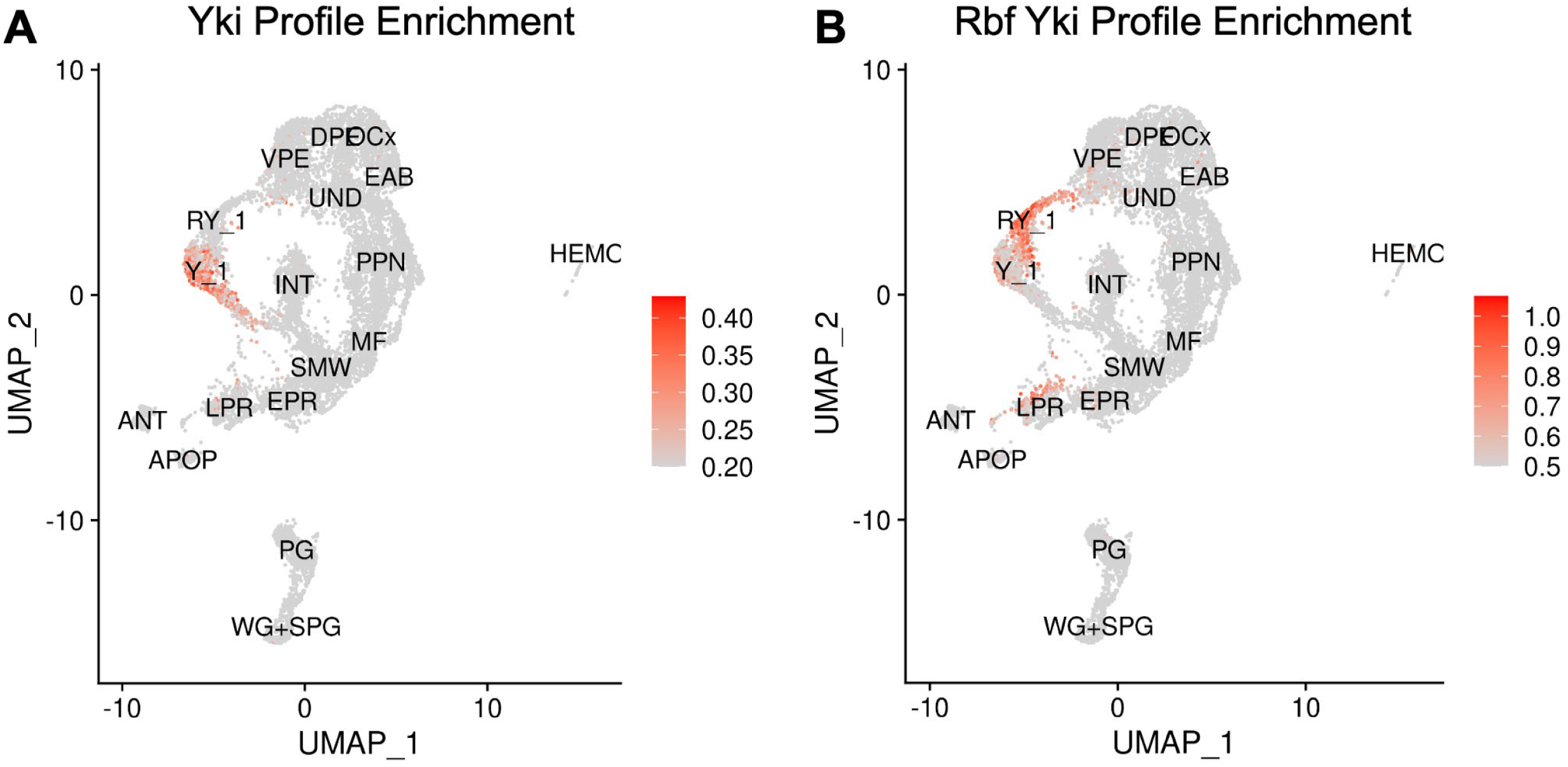
Y_1 and RY_1 have distinct and characteristic transcriptional signature. (a) A feature plot displaying expression of the Y_1 transcriptional profile. The Y_1 enrichment profile accurately and specifically captures Y_1 cells. (b) A feature plot displaying expression of the RY_1 transcriptional profile. The RY_1 enrichment profile accurately and specifically captures RY_1 cells.

**Figure S3.**
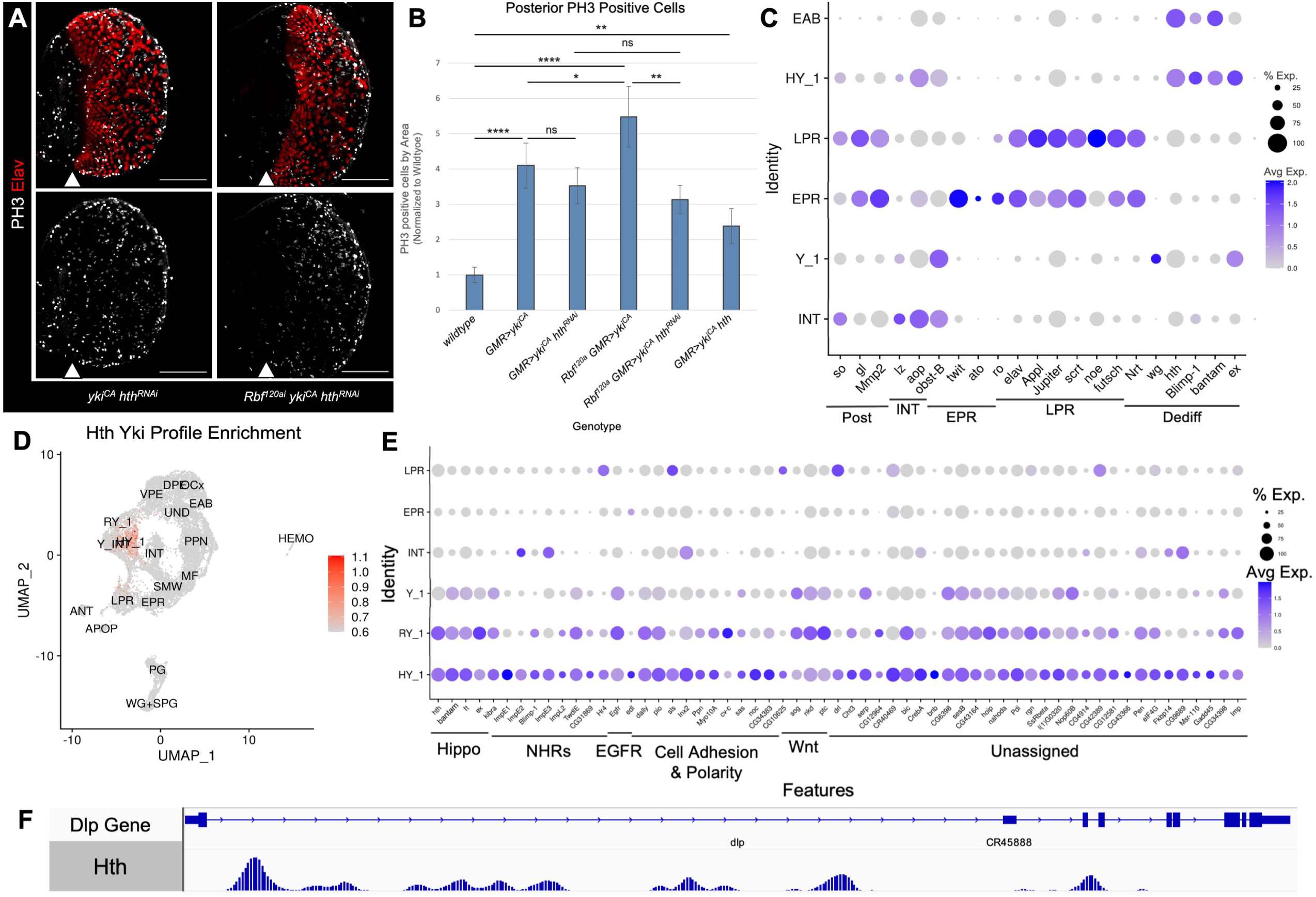
Hth/Yki activate proliferation and dedifferentiation programs. (a) Confocal images of third instar larval eye discs from and *+/+;GMR-Gal4/uas-hth^RNAi^;uas-yki^CA^ /+* and *Rbf^120a^/Y;GMR-Gal4/uas-hth^RNAi^;uas-yki^CA^ /+* animals immunostained for Elav (red) and PH3 (white). Hth knockdown tissue continues to proliferate. The MF is labeled with a white arrow. Scale bars are 100 μm (b) A bar graph displaying the number of mitotic, PH3-positive cells in the posterior compartment of each genotype. Hth knockdown does not significantly inhibit proliferation in *GMR>yki^CA^* tissues, which do not upregulate *Hth*. Hth knockdown in *Rbf^120a^ GMR>yki^CA^* eye discs reduces proliferation to *GMR>yki^CA^*levels. *GMR>hth yki^CA^* cells ectopically proliferate, though at levels lower than *GMR>yki^CA^*. (c) A Dot Plot of markers for posterior cells (Post), interommatidial cells (INT), early photoreceptor neurons (EPR), late photoreceptor neurons (LPR), RW_1 (Dediff), and the eye-antennal border (EAB). Expression is shown in corresponding cell populations, Y_1, and the *Hth yki^CA^* specific population HY_1. (d) A feature plot displaying expression of the HY_1 transcriptional profile. The HY_1 enrichment profile accurately and specifically captures HY_1 cells. (e) A Dot Plot of genes overexpressed in HY_1. Dedifferentiation models HY_1 and RY_1 significantly upregulate expression of NHRs, EGFR, and genes associated with cell adhesion and polarity. This is not seen in Y_1 cells. (f) ChIP-Seq data showing Hth occupancy at the *dlp* gene. *Dlp* contains several Hth peaks.

## Notes

### Competing Interest Statement

The authors have declared no competing interest.

